# Ribosome-induced mRNA pseudoknot interactions visualized by DMS MaP-Seq

**DOI:** 10.1101/2024.12.07.627321

**Authors:** Preston A. Kellenberger, Zhenwei Song, Xiao Heng, Peter V. Cornish

## Abstract

Examining the dynamicity of RNA structure has deepened our understanding of its vast biological functions. Perhaps the protein complex that encounters the most diverse landscape of RNA structure is the ribosome. In translation, the ribosome must linearize countless mRNA conformations for proper protein production. Some RNA structures, however, reliably make up sequences which hinder the ability of the ribosome to maintain its reading frame. The most well-studied of these structures is the RNA pseudoknot. Here, we present an approach utilizing dimethyl sulfate probing with mutational profiling and sequencing (DMS MaP-Seq) to precisely examine RNA unwinding. We employ the method to understand the unfolding of the Sugarcane Yellow Leaf Virus pseudoknot (ScYLV_PK_). Notably, we find that the helical junction is stabilized in the presence of the ribosome and is contingent upon hydrogen bonding at the 27^th^ residue of ScYLV_PK_. Additionally, it is demonstrated that the ribosome destabilizes wildtype ScYLV_PK_ in a manner independent of A/P-site occupancy. Together, these results establish DMS MaP-Seq as a sensitive tool for detecting ribosome-induced RNA conformational changes and reveal specific structural motifs that govern pseudoknot stability during translation.

## Introduction

In translation, the ribosome can only interpret single-stranded RNA to allow proper codon-tRNA alignment. However, complex intramolecular RNA secondary/tertiary structures are commonplace in transcripts. These structures, then, must be linearized before the bacterial ribosome is able to read their corresponding nucleotides. Central in aiding linearization of mRNA within bacteria is the ribosomal helicase center composed of proteins S3, S4, and S5 located within the 30S subunit [1]. Unlike DNA helicases such as DnaB, which are tasked only with unwinding the double helix conformation, S3, S4, and S5 must be able to unwind extensive combinations of RNA structure [2]. Fascinatingly, despite such vast modes of operation, it has been shown that these ribosomal helicase proteins do not require direct use of ATP or GTP [1]. Furthermore, little is understood about the underlying mechanical function of proteins S3, S4, and S5. These unique attributes of the ribosomal helicase center, compounded with the dynamicity of the ribosome, make ribosomal unwinding a process of which much is unknown.

-1 programmed ribosomal frameshifting (−1 PRF) is a phenomenon that resides centrally within the uncertainties of mRNA unwinding. Due to evolutionary pressures on their relatively small genome, many viruses utilize −1 PRF as a means to produce multiple proteins from a single coding mRNA [3]. In −1 PRF, the ribosome slips out of the original reading frame by shifting one nucleotide backwards, producing a different polypeptide. Three components are necessary for −1 PRF to occur: a slippery sequence (X XXY YYZ where X is any nucleotide, Y is A or U, and Z is A, U, or C), a downstream stable structure (ex. *dnaX* hairpin, HIV-1 hairpin, luteoviral pseudoknots), and a linker/spacer sequence positioned between the previous two elements [4–6]. It has been demonstrated that the length of the linker sequence (which typically ranges from 5nt-9nt) is vital for substantial rates of frameshifting [7]. Many have shown that the linker length needs to be short enough to provide tension between the ribosome and the downstream secondary structure, but not too short to where the ribosome has partially or completely unwound the structure before arriving at the slip site [8, 9]. Preliminary studies speculated that the thermostability of secondary structures found in frameshifting capable RNAs would provide resistance to the ribosome.

However, it is now known that the T_m_ of the mRNA has no direct correlation on frameshifting efficiency [10–12]. Therefore, the intricate unfolding patterns of these mRNA structures are fundamentally linked to their interaction with the ribosome, which is crucial for understanding how the ribosome processes mRNA and influences frameshifting efficiency.

Starting nearly three decades ago, a wide array of experimental approaches have been employed to understand the unwinding of mRNA pseudoknots. Initially, early thermodynamic studies, including differential scanning calorimetry (DSC) and UV–vis thermal denaturation, clarified global unfolding pathways but could not resolve nucleotide-level changes or 3D structures relevant to PRF [13–18]. X-ray crystallography and NMR experiments overcame these barriers through atomic resolution of many pseudoknots [19–27]. However, similar to thermodynamic studies, the effectiveness of these approaches is limited in the presence of the ribosome. Force measurements have also been used in unwinding experiments, measured through nanopores or tweezers. In ribosome-free contexts, tweezers revealed a hierarchy of denaturing interactions but lacked nucleotide precision and introduced extraneous 3′ forces [10, 12, 28–32]. Nanopores attempt to mimic the ribosomal mRNA entry tunnel by pulling mRNA through the channel while reading blocking signatures, indicating possible unfolding states [33]. Nanopores, however, present imprecise unfolding intermediates similar to that of tweezers. Additionally, all the above methods fail to consider how the complexity and dynamicity of the ribosome affect unwinding of RNA pseudoknots. In an effort to overcome this barrier, mutational experiments have speculated regions of the pseudoknots that may provide stability by analyzing relative frameshifting efficiencies [11, 34, 35].

Recently, tweezers experiments have been performed in the presence of the ribosome, providing a more accurate depiction of unwinding while still suffering from imprecision and extraneous 3′ forces [36–39].

Cryo-electron microscopy structures of early-stage ribosome/pseudoknot interaction have been released, but are only able to image well defined states, leading to difficulty in visualizing the heterogeneous partially-unwound RNA [40]. Probing techniques, such as RNases or dimethyl sulfate (DMS), permitted nucleotide-level precision in their results, but were traditionally limited in sensitivity [14, 41, 5, 42, 43]. Despite the benefits and limitations of these approaches, the need to examine ribosomal unwinding with nucleotide-level precision persists.

In a traditional DMS experiment, RNA secondary structure can be deduced *de novo*. DMS methylates A and C nucleotides at Watson-Crick positions (N^1^ and N^3^ respectively) and N^1^/N^7^ of G [44–47]. Although many factors contribute to the modification rate of a given nucleotide, Watson-Crick interactions and hydrogen bonding reliably reduce the probability of modification [48, 49]. Unlike classical DMS probing, which relies on gel analysis, the advent of DMS mutational profiling and sequencing (DMS MaP-Seq) allows for a greater degree of data precision and high reproducibility, enabled by next-generation sequencing of modified oligonucleotides [48, 50]. Thus, DMS MaP-Seq permits the identification of the relative strength of hydrogen bonding (bond flexibility) as a function of solvent accessibility [50, 51]. The amount of experimental freedom is another highlight for the use of DMS MaP-Seq. Whereas many approaches substitute the ribosome’s unwinding force, the simultaneous use of DMS MaP-Seq with the ribosome requires little additional effort beyond setting up ribosome/mRNA constructs. Furthermore, adding tRNAs and elongation factors in DMS MaP-Seq experiments allow for the investigation of ribosomal state as it pertains to mRNA probe reactivity [52]. In a 2009 study, Mazauric et. al used DMS probing in the presence/absence of the ribosome on the beet western yellow virus pseudoknot (BWYV_PK_) [42]. In these experiments, ribosomal unwinding of BWYV_PK_ was investigated by stalling ribosomes upstream (11-12 nucleotides from the P site) of the RNA structure. We sought to expand the scope of these experiments by utilizing DMS MaP-Seq and by leveraging multiple ribosomal states. In the attempts to elucidate how these mRNAs are unwound, we have chosen the Sugarcane Yellow Leaf Virus (ScYLV) pseudoknot as the basis of our experimentation. The ScYLV pseudoknot (ScYLV_PK_) has been well characterized in multiple studies, and its structure has been solved through NMR [23, 24]. Additionally, the wild-type ScYLV_PK_ contains a larger frameshifting efficiency than that of BWYV despite both originating from luteoviridae [24, 42]. However, as is the case with ScYLV_PK_ and all other mRNA structures found within frameshifting cassettes, little is known about how the RNA is unwound in the process of translation.

## Results

### Probing folded ScYLV_PK_ yields distinct mutational profile

In this study, we sought to examine the destabilization of ScYLV_PK_ with single-nucleotide resolution using DMS MaP-Seq. Traditional DMS MaP-Seq experiments aim to infer RNA secondary structure *de novo* using DMS-derived reactivity profiles. Instead, we apply DMS MaP-Seq to detect localized changes in nucleotide flexibility with an informed understanding provided by the solution structure of ScYLV_PK_ (Figure 1A) [23]. To achieve this, we constructed a 117-nt. mRNA that contains a ribosomal binding site (RBS), AUG and UUU codons for tRNA binding, variable linker sequence, ScYLV_PK_, and a trailing sequence (Figure 1B). The incorporation of the RBS permits the ribosome to bind upstream of ScYLV_PK_ in a stalled state, enabling *in-vitro* probing of the mRNA with the ribosomal P site positioned at singular codon. There remains uncertainty surrounding the existence of non-canonical rotational states of −1 frameshifted ribosomes, which may interact with ScYLV_PK_ in different manner than non-frameshifted ribosomes. Replacing the native slip site with the RBS permits the experimenter to isolate the effects of unwinding without an unknown population of ribosomes entering the −1 frame. Adding the unstructured trailing sequence supplements mutational normalization with more points of reference (See Experimental Procedures).

**Figure 1:**
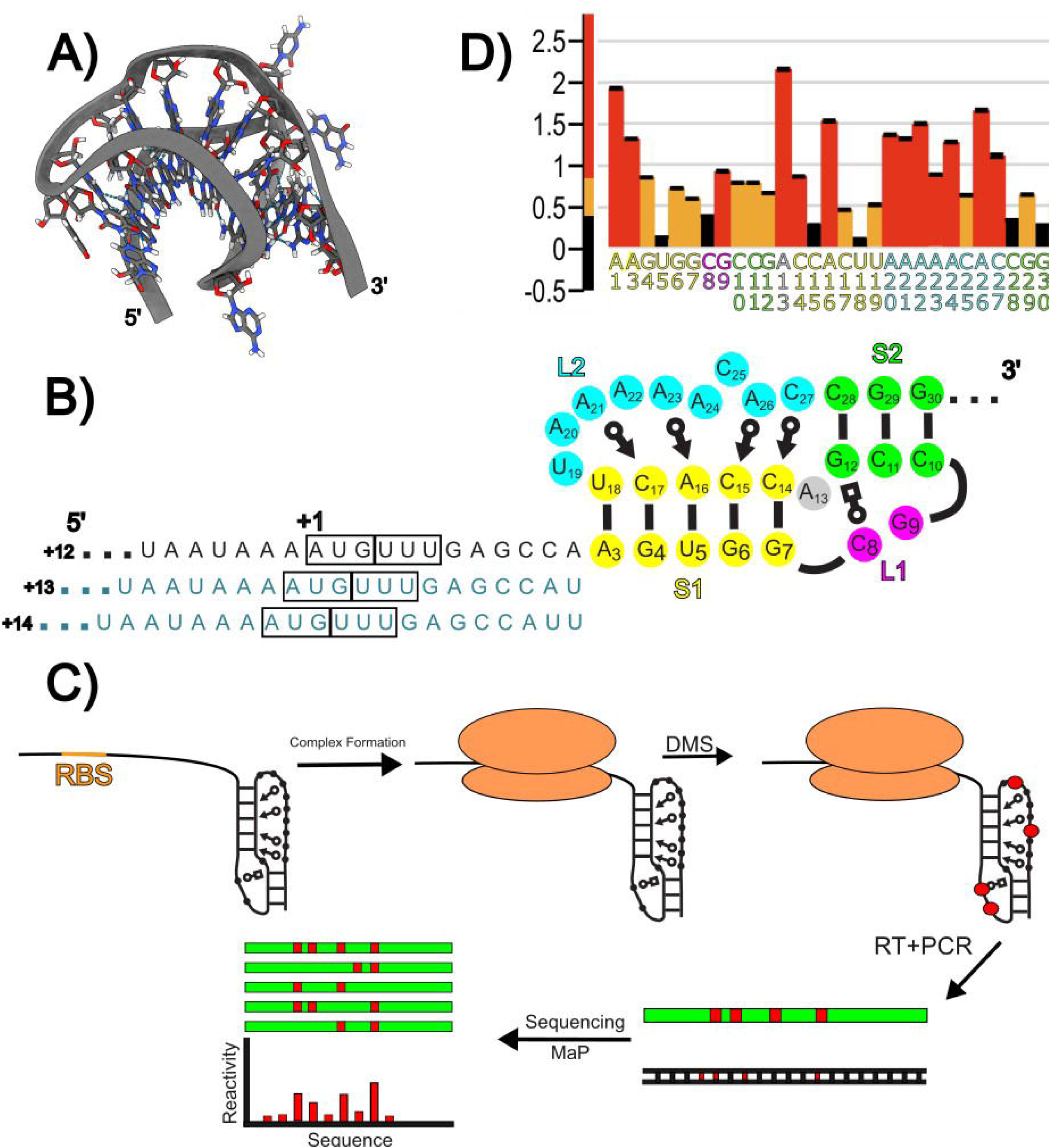
Workflow of in-vitro ribosome mRNA DMS probing experiments. A) NMR structure of ScYLV_PK_ PDB: 1YG4. B) 2D representation of experimental RNA containing ScYLV_PK_. Boxes surround location of ribosomal P site (AUG) and A site (UUU). “+1” above 5’ P-site nucleotide is shown. +13 and +14 variable linker sequences are pictured in blue. Symbols within ScYLV_PK_ notate non-Watson-Crick interactions as identified by Leontis et. al [81]. C) Experimental outline of DMS MaP-Seq with stalled ribosome-mRNA complexes. Red dots symbolize locations of RNA modification. D) A DMS reactivity profile of ScYLVPK at single-nucleotide resolution in the absence of the ribosome.

In its native state, folded ScYLV_PK_ contains two loops (L1/L2) and two stems (S1/S2) with three nucleotides that have no binding partners (G9/C25/A13) (Figure 1B). Before probing ScYLV_PK_, we speculated that L1/L2 should exhibit increased DMS reactivity as compared to S1/S2 due to decreased steric impedance. To probe ScYLV_PK_, we expose the RNA to DMS over a period of 7 minutes (See Experimental Procedures) (Figure 1C). Reverse transcription of the RNA introduces mutations at methylated nucleotides. The addition of sequencing adapters to the 5’ and 3’ end of the cDNA enables read collection (∼1 million reads per experiment). After aligning these reads, the mutational frequency at each nucleotide is determined. The resulting mutational histogram (referred to as a mutational profile) indicates probe-accessibility on a nucleotide-level basis. Probing ScYLV_PK_ in the absence of the ribosome yields a mutational profile where S1/S2 nucleotides are significantly less reactive than L1/L2 nucleotides (Figure 1D). Although some pseudoknots do not show reactivity at L2 adenosines[53] as seen in our Figure 1D, it is important to note that the N^1^ of ScYLV_PK_ L2 adenosines (21, 23, and 26) are involved in unusual minor-groove base triples in which the N^1^ of adenosine can form a hydrogen bond with 2’ OH in S1 residues [23]. This interaction is weaker than a standard base triple interaction as seen in C8+-G12-C28. Therefore, the reactivity for L2 residues are increased relative to the reactivity for C8, G12, and C28 in the ribosome absent state (Figure 1D) [23]. Indeed, probing of a highly similar luteoviral pseudoknot, BWYV_PK_, has been shown to exhibit strong DMS reactivity at L2 adenosines [42]. Consistent with the NMR structure of ScYLV_PK_, strong DMS reactivities at positions G9 and A13 are also observed (Figure 1D). In addition, the same refolding protocol used to determine the structure of ScYLV_PK_ was followed as in our previous studies (See Experimental Procedures) [23, 24]. Taken together, these results indicate probing of properly folded ScYLV_PK_.

### The presence of the ribosome changes pseudoknot flexibility

One critical element within frameshifting is the distance between the ribosome-occupied slip site and the downstream RNA structure. The distance parameter is finely tuned by the linker (or spacer) sequence [9]. Linker sequences place the RNA structure at the helicase center of the ribosome, which resides +11nt from the 5’ residue in the bacterial ribosomal P site [1, 7]. Arginine- and lysine-rich helicase proteins (S3, S4, and S5) occupy the helicase center and are thought to interact with the phosphodiester backbone of the transcript [1]. A select few studies have attempted to examine ribosomal unwinding of RNA, but, due to technique limitations, are unable to observe nucleotide-level changes in RNA structure [36–38]. Additionally, it has been observed that −1 frameshifting efficiency drops sharply when the downstream structural element is positioned further than +13 from the ribosomal P site [9]. Considering these parameters, we constructed +12, +13, and +14 ScYLV RNAs to investigate changes in ScYLV_PK_ flexibility. The RNAs are identical except for the addition of uracil residue(s) preceding A3 (Figure 1B) of the pseudoknot. In the absence of the ribosome, we find that the +12, +13, and +14 RNAs show minor change in their respective mutational profiles when compared against each other (Pearson; 0.8 < ρ < 0.82), indicating modification of linker length does not disrupt ScYLV_PK_ stability (Figure S1).

In addition to studying ScYLV_PK_ with respect to ribosomal distance, we also sought to determine the effect of P/A site occupancy. It has been found that the addition of A – site tRNA causes the ribosomal entry tunnel to constrict 1-3 Å, an effect which may play a role in both intersubunit rotation and unwinding [54]. As reported from previous smFRET data, the ribosomal state can be manipulated via the addition of tRNAs, elongation factors, GTP, and other various reaction components [52, 55]. Such perturbations can shift equilibria between the classical and hybrid states, which define the ratcheting motion of the translating ribosome. In previous fluorescence studies, we have identified an additional state of hyper rotation, characterized by a 22° counter-clockwise rotation between large and small subunits [55]. The hyper rotated state appears to form when the ribosome is within close proximity to a downstream nucleic acid structure [56]. Since DMS MaP-Seq cannot provide information on the rotational states of individual ribosomes, we will be referring to our experimental conditions with the following notation: acylated P – site tRNA (aP-tRNA) (traditionally shifts equilibrium to the classical state), deacylated P – site tRNA (daP-tRNA) (traditionally shifts equilibrium to the hybrid state), and daP-tRNA with acylated A – site tRNA to construct the pre-translocation state (PRE). In the experiments detailed below, each variable linker mRNA (+12, +13, +14) was subject to ribosomes in the aP-tRNA, daP-tRNA, and PRE states.

To capture changes in pseudoknot DMS reactivity, we used a two-phase analysis, permitting comparison of ribosome bound and absent states of ScYLV_PK_. First, we applied the widely-utilized method of 2/8 normalization to limit the effect of fluctuations in universal modification rate [57–61]. Specifically, 2/8 normalization adjusts the global rate of modification by considering DMS reactivity in the unstructured 3’ trailing sequence, which is assumed to not interact with the ribosome and readily modified by DMS (See Experimental Procedures). Secondly, to detect small but significant changes in DMS reactivity, we calculated the z-score of the mutational frequency difference for each ribosome-bound condition relative to the no-ribosome control. The final normalized z-scores permit direct comparison between experimental conditions. Subsequent mapping of the mutational difference z-scores generated color maps, which were imposed on a 2D projection of ScYLV_PK_ (Figure 2).

**Figure 2:**
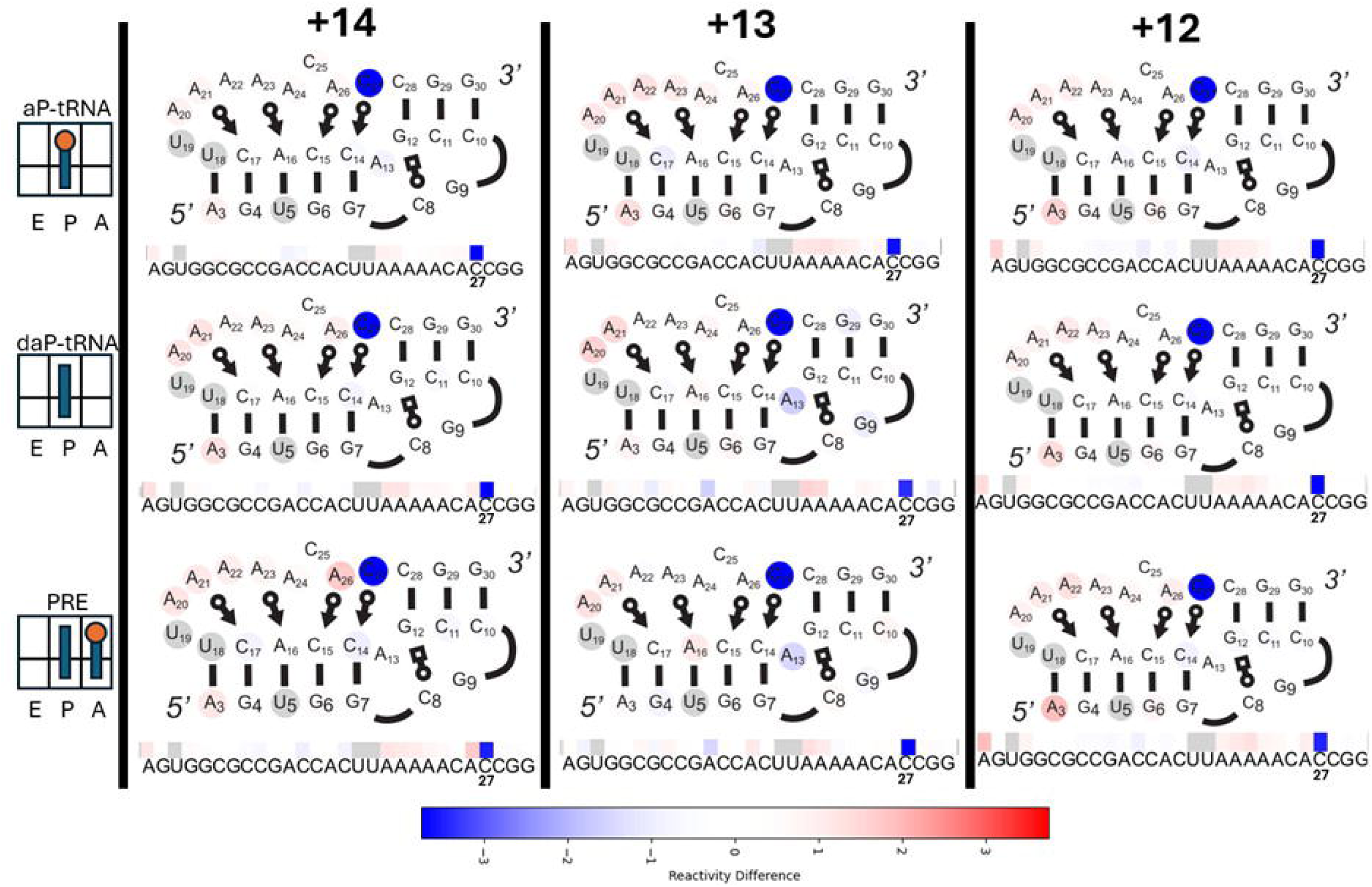
Changes in ScYLV DMS reactivity in the presence of the ribosome. A display of z-score changes (with respect to no ribosome) across +12, +13, and +14 WT ScYLV RNAs under aP-tRNA, daP-tRNA, and PRE ribosomal states. Z-score heat maps are displayed beneath 2-dimensional (2D) representations of ScYLV_PK_ along with the RNA sequence. Heat map coloration is applied to the 2D projection of ScYLV_PK_. The 27^th^ position is labeled under each experiment for ease of identification.

Performing these experiments, we hypothesized that ribosomal state and position would greatly affect ScYLV_PK_ stability. Contrary to our expectations, we observe that the reactivity changes induced by the ribosome are nearly identical in every tested condition (Figure 2). The reproduced pattern of flexibility change seems to follow sub-structural trends, where reactivity increases near the ribosomal footprint and decreases in distal regions, further from the ribosome. Specifically, L2 adenosines (A20-A24) moderately increase in flexibility, exhibiting a gradient in magnitude of change towards the 3’ end of L2. Additionally, the first base pair in S1 (A3 – U18) shows increased mutational rate as compared to a no-ribosome control. Mazauric et al. also observed the first S1 WC pair increasing in DMS reactivity in their probing of the BWYV_PK_ [42]. Simultaneously, decreases in reactivity are observed in regions furthest away from the ribosomal footprint (G9, A13, C14, C27, and G29). The rigidity found in G9, A13, C14, C27, and G29 seem to be localized around an area of ScYLV_PK_ known as the helical junction. The helical junction is defined by the coaxial stacking of S1 and S2 and has been speculated to provide resistance to the ribosome in frameshifting [10, 12, 62–64]. Most significant among any flexibility change is C27, exhibiting a sharp decrease in DMS reactivity as compared to the rest of the molecule across all experiments (z_WT_ = −4.01 ± 0.34).

#### Mutation C27A changes pseudoknot flexibility in the presence of the ribosome

Previously, we characterized a point mutation, C27A, that decreases the frameshifting efficiency of ScYLV_PK_ (15% to 2%) while maintaining its 3D conformation [23, 24]. Subsequent thermodynamic experiments examined hydrogen bonding networks along the helical junction of ScYLV_PK_. We concluded that the hydrogen bonds formed by C27 were favorably coupled (δ_AB_^WT^ = −0.7 kcal/mol) while hydrogen bonds formed by A27 were not (δ_AB_^C27A^ = 0.9 kcal/mol) [65]. Strikingly, our wildtype ScYLV_PK_ DMS MaP-Seq results indicate that C27 undergoes the most significant change in hydrogen bond flexibility out of the entire molecule (Figure 2). Considering all of the above, we sought to examine how incorporation of adenosine in position 27 would affect ScYLV_PK_ stability by comparing z-scores of reactivity in the presence and absence of the ribosome, hypothesizing that C27A would decrease global pseudoknot stability. After verifying that the mutation was incorporated, we constructed +12, +13, and +14 mutant RNAs and subjected them to the same ribosomal states as detailed previously. RNAs were refolded as outlined in our past reports (See Experimental Procedures), and, additionally, in the absence of the ribosome, little reactivity profile differences between C27A and WT RNAs were observed (Pearson; ρ = 0.83) [23, 24]. Given these similarities, we next asked whether ribosome binding changes the destabilization of C27A ScYLV_PK_ as compared to WT ScYLV_PK_.

Consistent with our hypothesis, incorporation of C27A results in a pronounced increase in DMS reactivity at position 27 (z_C27A_ = 4.46 ± 0.54) across the majority of experimental conditions (Figure 3A depicts average change across all 9 conditions), representing an 8.5 z-score swing relative to wild type (z_WT_ = –4.01 ± 0.34). The extreme shift corroborates the importance of the 27^th^ position in determining frameshifting efficiency for ScYLV_PK_. Notably, even under these destabilizing conditions, helical junction residues G9 and A13 remain rigid. In contrast, a subset of experiments, including the +12 aP-tRNA, +14 daP-tRNA, and +14 PRE conditions, display a markedly different pattern resembling the wild-type experiments (See “Ribosomal state changes pseudoknot flexibility”). In these instances, A27 shows a substantial decrease in flexibility comparable to that of the native pseudoknot (z_WT_ = –3.00 ± 0.857, z_C27A_ = –4.01 ± 0.34). These experiments, like wild type, exhibit increased flexibility at L2 adenosines A20–A24 (Figures 2 and 3). Another interesting finding is that helical junction flexibility seems to be, to some degree, independent of A27. More specifically, G9 and A13 maintain decreased z-scores regardless of if A27 is highly flexible or highly rigid (Figure 3B). Taken together, these results indicate that the mutation C27A destabilizes the 27^th^ position as compared to the native sequence in the presence of the ribosome.

**Figure 3:**
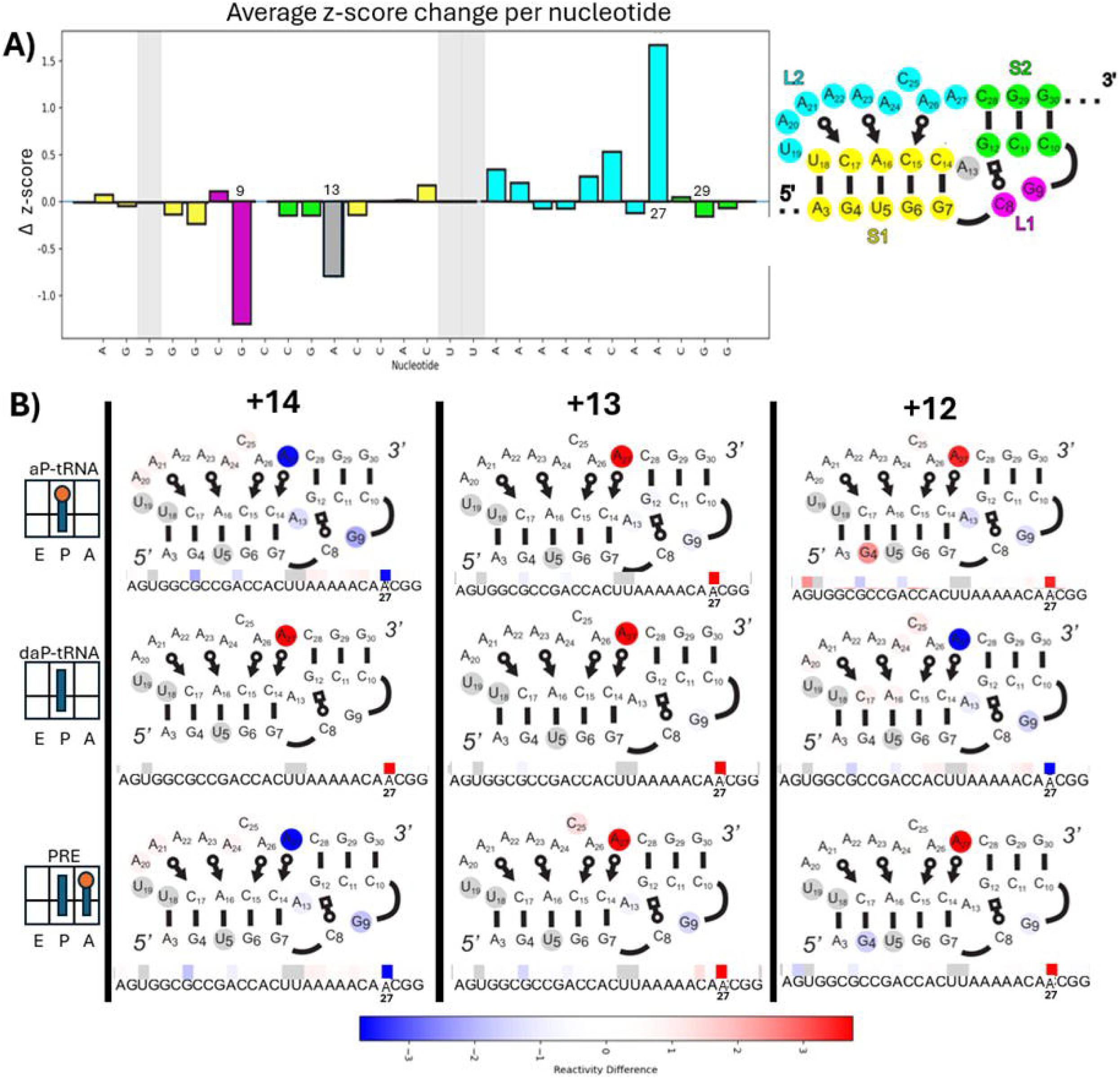
Changes in ScYLV C27A DMS reactivity in the presence of the ribosome. A) A bar graph depicting the average change in z-score across all experimental constructs (aP-tRNA, daP-tRNA, PRE) and linker lengths (+12, +13, +14) in C27A ScYLV RNA. B) A display of z-score changes across +12, +13, and +14 C27A ScYLV RNAs under aP-tRNA, daP-tRNA, and PRE ribosomal states. Z-score heat maps are displayed beneath 2-dimensional (2D) representations of ScYLV_PK_ along with the RNA sequence. Heat map coloration is applied to the 2D projection of ScYLV_PK_. The 27^th^ position is labeled under each experiment for ease of identification.

### N1 guanosine mutations reveal coordinated helical junction rigidity

Recently, it has been reported that G selectively mutates dependent upon N^7^ (G to A) or N^1^ (G to C/U) DMS modification [45]. Thus, isolating these mutations would allow delineation of WC/Hoogsteen face modification. In both the wildtype and C27A RNAs it appears as if the most significant changes in flexibility occur near the helical junction. Within the helical junction, G12 appears in non-canonical base pairing with C28 and C8+, a naturally protonated cytosine (Figure 4A). Due to the unique bonding present between G12, C8+, and C28, we sought to determine if any changes occurred within N^7^ and N^1^ modification rates. After handling G to A and G to C/U mutations we found that the rate of G12 N^7^ modification does not significantly change across any condition, indicating no observable change in the Hoogsteen face. In contrast, while slight or no significant change occurs between G12 N^7^ modification across all ribosomal states and RNAs, G12 N^1^ modification within the daP-tRNA state on C27A RNA is reduced, on average, by factor of 0.18 (Figure 4B). Intriguingly, G12 is constitutively rigid despite being adjacent to A27, which both sharply increases (+13 and +14) and decreases in modification rate (+12) (Figure 3B). Similarly, we found that N^1^ modification within C27A RNAs in daP-tRNA states decrease significantly in G6, G7, and G9 (Supporting Information). Taken together with z-score decreases present in A13 (Figure 3), these results indicate that nucleotides G6, G7, G9, A13, and G12 act in coordination across one or more states, increasing rigidity in daP-tRNA state within C27A RNA.

**Figure 4:**
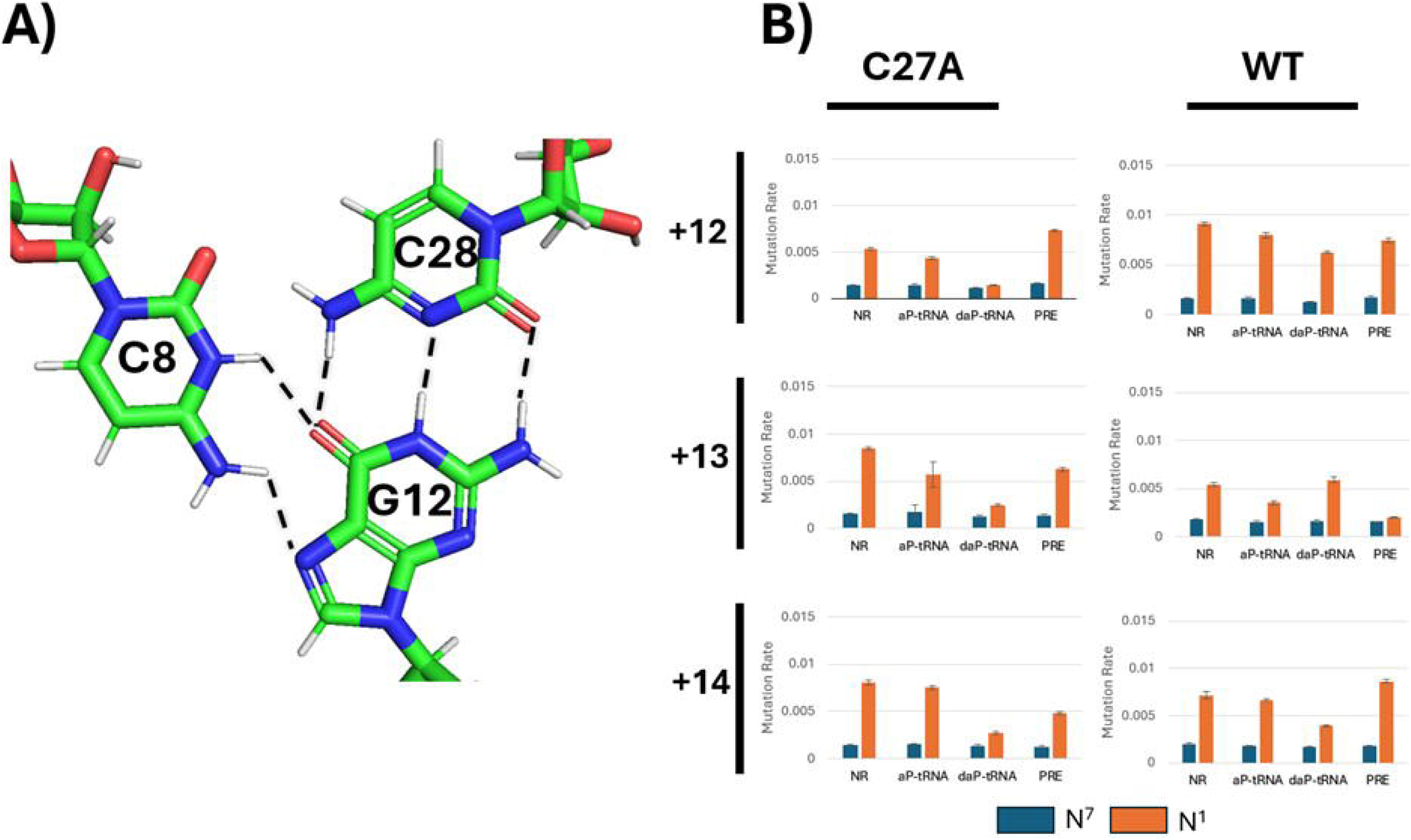
N1 and N7 modification perturbations within C8+-G12-C28 triple base pair. A) Structural view C8+-G12-C28 triple base pair (PDB: 1YG4). B) Bar graphs of N^7^ (blue) and N^1^ (orange) mutation rates of G12 in +12, +13, and +14 WT and C27A RNAs across all ribosomal states (daP-tRNA, aP-tRNA, PRE) for G12. Error bars represent 95% Wald confidence interval.

The ratio of N^7^:N^1^ mutations across the wild type +12 and +14 states were found to not change significantly across ribosomal construct for most guanines present in the pseudoknot (Supporting Information). The +13 wild type RNA, however, exhibited decreased N^1^ modification in G6, G7, G9, A13, and G12 when the ribosome was in PRE (Figure 4, Supporting Information). This observation is similar to the N^1^ labeling pattern observed within the daP-tRNA C27A experiments and may suggest a shared meta-stable state of ScYLV_PK_. It is worth noting that helical junction rigidity is, to an observable degree, somewhat independent of the flexibility of the 27^th^ residue of ScYLV_PK_, appearing both reactive and non-reactive while G6, G7, G9, A13, and G12 maintain rigidity. Together, our results demonstrate that DMS MaP-Seq can sensitively detect subtle yet significant perturbations in nucleotide flexibility when applied to a known RNA structure. These perturbations revealed distinct conformational states under experimental conditions that would otherwise be difficult to resolve using conventional structural methods.

## Discussion

In this study, we sought to characterize ribosome-induced flexibility changes within ScYLV_PK_ by applying DMS MaP-Seq to an RNA with a previously determined structure. Through the contrast of these changes, we aimed to identify the dynamic interactions of RNA pseudoknots that influence frameshifting efficiency. By probing wild type and mutant forms of ScYLV_PK_ with respect to varying ribosomal states we observed significant and precise changes within ScYLV_PK_ stability, with most alterations occurring proximal to the helical junction.

One of the main objectives in our investigation was to expand upon the works of Mazauric et al. [42]. Previously, in their 2009 report, DMS footprinting was performed on BWYV_PK_ in the presence and absence of the ribosome. The traditional use of DMS footprinting relies on the chemically induced cleavage of methylated ribonucleotides that are imaged through polyacrylamide gel electrophoresis [66]. As motivation, we posited that ribosome-induced mRNA stability changes are both dynamic and subtle and therefore would benefit from the enhanced precision granted by DMS MaP-Seq. Beyond the technique, we altered the experimental setup to capture a wider range of ribosomal dynamics, expanding Mazauric et al.’s PRE-only analysis of BWYV_PK_ to also include daP-tRNA and aP-tRNA states.

Additionally, our experiments focus on RNAs lacking a slip site. We believe that by removing the slip site, we can remove heterogeneity arising from frameshifted and non-frameshifted populations; thus, more accurately isolating the unwinding effects of the ribosome. The results of our experiments share some similarities and a few prominent differences. In agreement with the 2009 study, our data indicate that the ribosome stabilizes tertiary interactions surrounding the helical junction at the back of S1 and L1 and destabilizes WC pairing near the 5’ end of S1 (Figure 2). Conversely, while Mazauric et al. observed rigidity of the entire L2 region, we have found that most of L2 exhibits slight destabilization (Figure 2). The differences between our results may arise from the newfound sensitivity of MaP-Seq; however, it is also possible that BWYV_PK_ is destabilized entirely differently by the ribosome, resulting in the reduced frameshifting efficiency as compared to ScYLV_PK_ ^[23,42]^.

The most intriguing result of our study is the observation of two reproducible DMS reactivity-change signatures within ScYLV_PK_. The primary distinguishing element between these two signatures is position 27, either exhibiting substantial stabilization or destabilization (Figure 5A). Signature I is characterized by strong stabilization of the 27^th^ position with concomitant rigidity in the helical junction (G9, A13, position 27, and G29), increased flexibility in nucleotides L2 (A20-24), and increased flexibility in 5’ WC pairing (A3). Interestingly, the flexibility in L2 follows a stepwise regression in signal moving downstream (Figures 2 and 5A). It is possible that this is a result of the force of tension being propagated through L2, originating from the ribosome in an attempt to unwind ScYLV_PK_. Conversely, signature II is characterized primarily by enhanced flexibility at position 27 and rigidity of the helical junction, although to a lesser degree than signature I (G9 and A13). In probing the WT ScYLV_PK_ at varying ribosomal distances and states, we observed signature I in every scenario (Figure 2). Such a finding is surprising, as the frameshifting efficiency of an RNA is highly dependent upon its linker/spacer distance. Our data support that linker/spacer lengths may promote frameshifting in a manner agnostic to downstream structural RNA unwinding, contradicting hypothesis such as the torsional restraint model [67]. In contrast, introduction of mutation C27A resulted in visualization of signature II in the majority of tested conditions (Figure 3).

**Figure 5:**
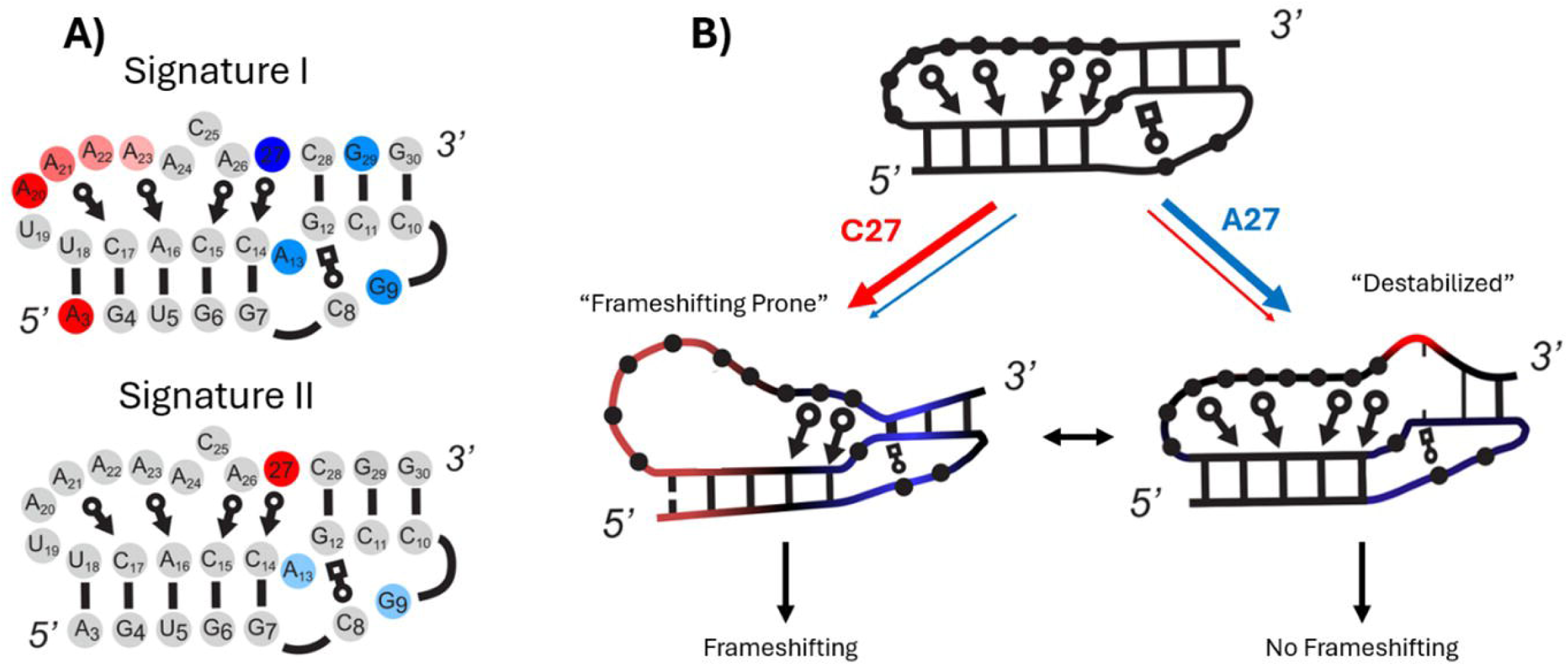
Frameshifting efficiency changes explained by DMS signatures. A) Representations of DMS reactivity change signatures I and II presented on 2D view of ScYLV_PK_. B) Proposed pathway of stability changes induced by the ribosome upon ScYLV_PK_ from a basal to “frameshifting prone” and “destabilized” states.

Interestingly, WT BWYV_PK_ contains adenosine at the analogous position 27 of ScYLV_PK_. Despite this similarity, signature II does not seem to appear in Mazauric et al.’s probing BWYV_PK_, supporting that ribosomal interaction with BWYV_PK_ is different than that of ScYLV_PK_. As mentioned previously, we have found that the mutation C27A maintains a structure nearly identical to that of WT ScYLV_PK_ while decreasing frameshifting efficiency nearly 4-fold [24]. Consecutive experimentation with pairwise coupling showed that C27A thermodynamically destabilizes the helical junction [68]. The majority presence of signature II within the concurrent C27A data further suggests that non-covalent interactions in the 27^th^ position are critical in ribosome-induced ScYLV_PK_ dynamicity, possibly accounting for the significant decrease in frameshifting efficiency as compared to wild type RNA.

The flexibility changes observed in the preceding experiments provide an explanation as to which transient intramolecular RNA hydrogen bonds provide mechanical resistance against the ribosome. The foundation of this proposition lays within the enhanced rigidity demonstrated within the helical junction, a region speculated to be instrumental in determining frameshifting efficiency across many studies [10, 12, 30, 35, 69]. As shown in our N^1^ guanosine analysis, G residues proximal to the helical junction (G6, G7, G9, G12, and G29) become rigid in the +13 WT PRE state and all daP-tRNA C27A RNAs compared to no-ribosome states (Figure 4B, Supporting Information). Additionally, residues G9 and A13, which reside near the helical junction and contain no binding partners, are found to decrease in DMS reactivity, possibly indicating that these residues become buried within the helical junction. In both our z-score and G mutation analysis, the stability of the helical junction exists regardless of the flexibility state of the 27^th^ position; however, the guiding principles as to why specific ribosomal states and distances elicit helical junction rigidity remain unclear. We believe the strong stabilization of position 27 and residues within the coaxial stacking of S1/S2 provide resistance to the ribosome, which may facilitate entry into the long-paused “frameshifting prone” state as observed in many studies [4, 29, 70–75]. In addition, we propose that that signature II represents a destabilized state of ScYLV_PK_ along the unfolding pathway, whereby flexibility at the 27^th^ position weakens the helical junction. Based on our findings, we present a stability pathway of ScYLV_PK_, summarized in Figure 5B.

Previous optical tweezers studies have attempted to define unfolding pathways of mRNA pseudoknots in order to better inform a mechanism of −1 PRF. Many tweezers experiments suggested the importance of triplex interactions in accordance with unfolding resistance [30, 35, 76]. However, universal mechanical unfolding of RNA pseudoknots with tweezers is not correlated with frameshifting efficiency [30, 76]. The selective stabilization of ScYLV_PK_ minor-groove triplex interactions within our data explains the uncoupling of frameshifting efficiency to universal mechanical unfolding. However, the importance of individual triplex interactions are likely dependent upon the specific frameshifting sequence, as some studies have found only major-groove interactions to promote frameshifting [30, 77]. Additionally, many tweezers experiments present intermediate structures where entire stems/loops are completely unwound in one step [29–31]; however, the extent to which these results inform a frameshifting mechanism is inherently limited. It is unlikely that frameshifting occurs when an entire stem or loop is unwound. Rather, it is more probable that the intricate process of destabilizing pseudoknot substructures promotes a −1 PRF prone state. Within frameshifting constructs, the roles of the slippery site and linker sequence have been defined and well supported. Conversely, the mechanistic understanding of structural elements in frameshifting systems has been elusive. We believe that DMS Map-Seq provides a promising tool to investigate precise interactional changes in frameshifting RNAs arising from ribosomal presence.

With the simultaneous use of DMS MaP-Seq with ribosome/mRNA constructs, we have leveraged the precision of the technique to elucidate how unique interactions within ScYLV_PK_ change due to the ribosome. We have found that hydrogen bonding within the helical junction creates substructural rigidity within ScYLV_PK_ when in the presence of the ribosome. Furthermore, helical junction rigidity dissipates when position 27 is destabilized by the ribosome. Previous attempts to understand ribosome/mRNA were limited precision and application in the presence of the ribosome. The use of DMS MaP-Seq addresses both of these constraints simultaneously while remaining simple to integrate within traditional in-vitro ribosome experiments. Despite the broad experimental approaches applied, the role of RNA pseudoknots in FSS is still not fully understood. Ribosomal frameshifts may affect how the ribosome interacts with the structural element. Re-folding of a pseudoknot may place it in a non-native conformation after a ribosome has translated through. Additionally, the RNA pseudoknots may influence the structural dynamics of the ribosome. Further investigation is required to answer much of the above items in order to paint a clearer picture of ribosomal frameshifting.

## Experimental Procedures

### Construction and Purification of RNA and Ribosomes

Construction of a pUC19 vector plasmid containing the ScYLV sequence was carried out via sequence and ligation independent cloning (SLIC). Purification and digestion of the plasmid with EcoRV (NEB, R0195S) produced a linear DNA fragment containing the T7 promoter and ScYLV sequence (Supporting Information). The DNA was then transcribed into mRNA via T7 RNA polymerase. Further isolation of the RNA was done through crush and soak purification [78]. Tight couple ribosomes (E. *coli,* MRE600) were purified as described by Staehelin and Maglott [79].

### tRNA Preparation

tRNA^fMet^ and tRNA^Phe^ were purchased from MP Biomedicals and Sigma respectively.

Charging of the tRNA was done by the following: 4X methionine charging buffer (50mM Tris HCL pH 7.9, 150mM KCl, 7mM MgCl2, 0.1mM EDTA, 2.5mM ATP, 1mM DTT) or 4X phenylalanine charging buffer (100mM Tris HCL pH 7.9, 50mM KCl, 0.5mM EDTA, 2.5mM ATP, 1mM DTT) were mixed with tRNA (20µM final), phenylalanine (100µM final) or N10-formyl tetrahydrofolate (300µM final), 4X S100-tRNA extract (0.0125µL per pmol tRNA), and were incubated at 37°C for 10 minutes. The reactions were quenched by the addition of 0.1 volumes of 3M NaOAc pH 5.2. Following phenol chloroform extraction, the samples were resuspended in MilliQ water to a final concentration of 40 µM. The samples were then run on a P6 column preequilibrated in 10 mM KOAC pH 6.1.

To acetylate Phe-tRNA^Phe^, acetic anhydride was added to the sample. Following the addition of acetic anhydride, the sample was put on ice for 1 hour. The acetic anhydride and resting process was repeated an additional time. Then, following the addition of 2 volumes of 100% ethanol, the sample was transferred to a dry ice ethanol bath for 30 minutes. The tRNA was then spun down and run through a G25 spin column equilibrated in 1mM KOAc pH 5.3.

### DMS Exposure

To ensure the RNA would assume the correct pseudoknot conformation, 10nM of ScYLV RNA was prepared via refolding in accordance with our previous studies of ScYLV [23, 24]. For this, the RNA was heated to 60°C for 5-10 minutes and then cooled on ice. 5X buffer was then added to the solution (250mM Tris pH 7.5, 500nM NH4Cl, 100mM MgCl2). Tight couple purified ribosomes (E. *coli,* MRE600) were added to the mix to a final concentration of 20nM, and samples were incubated at 37°C for 5 minutes. tRNA (final concentration 30nM) was then added to a final reaction volume of 10µL, mixed, and incubated for 20-30 minutes at 37°C.

After incubation, 15µL of 1X ribosomal buffer (50mM Tris pH 7.5, 100mM NH4Cl, 20mM MgCl2, 6mM BME) was added to the reaction, followed by the addition of 12µL of 1M bicine. Each reaction was then brought 37°C. 4µL of 1.7M DMS-Ethanol was added to the reaction tube and was incubated at 37°C for 7 minutes. The reaction was quenched with the addition of 200µL of BME Stop Solution (BME, 2ng carrier RNA).

### RNA Purification and Library Generation

To precipitate the RNA, 960ul of 100% ethanol and 1.5 µL glycogen was added to each reaction. The RNA was then precipitated at −80°C, spun and washed with 100% ethanol three times, and resuspended in 50uL of RNAase free water. The resulting material was reverse transcribed (M-MLV RT) and purified as described by Smola, M. et al. (NEB Monarch DNA Cleanup Kit) [80]. Two rounds of PCR were used to tag each reaction with sequencing barcodes (Illumina, Supporting Information). After verifying the length of the product on a gel, the samples were pooled and sequenced (Illumina NovaSeq 6000, 200M PE reads). The resulting data were analyzed via Shapemapper2 to obtain mutational profiles (Supporting Information).

### Handling Mutational Rates

Raw non-normalized mutational outputs were generated via shapemapper2 and handled separately. The 2-step data handling procedure first normalizes mutational rates by 2-8% normalization, then calculates z-scores of reactivity difference for each experimental condition. For 2-8% normalization, we rank the top 10% of highly reactive nucleotides in the unstructured trailing sequence. The most reactive 2% are discarded, and the remaining 8% of reactivity are averaged to create a scaling factor which is applied to the rest of the molecule. To calculate the z-score of reactivity change, normalized experimental mutational rates are subtracted from a baseline in the absence of the ribosome. Then, the standard deviation of rate change is computed in each construct, and z-scores are determined.

## Data availability

Shapemapper2 analysis and raw sequencing data is available upon request (Preston A. Kellenberger).

## Supporting Information

This article contains supporting information.

## Supporting information

Target profiles 1

Target profiles 2

Target profiles 3

Target profiles 4

Target profiles 5

Target profiles 6

Target profiles 7

Target profiles 8

Target profiles 9

Target profiles 10

Target profiles 11

Target profiles 12

Target profiles 13

Target profiles 14

Target profiles 15

Target profiles 16

Target profiles 17

Target profiles 18

Target profiles 19

Target profiles 20

Target profiles 21

Target profiles 22

Target profiles 23

Target profiles 24

## Acknowledgements

We thank the University of Kansas Medical Center – Genomics Core for generating the sequence data sets. The Genomics Core is supported by the Kansas Intellectual and Developmental Disabilities Research Center (NIH U54 HD 090216), the Molecular Regulation of Cell Development and Differentiation – COBRE (P30 GM122731), The NIH S10 High End Instrumentation Grant (NIH S10OD021743) and the Frontiers CTSA Grant (UL1TR002366).

## Funding and additional information

Mizzou Forward Undergraduate Research Training Grant, University of Missouri Agricultural Institute, and NSF: #2122902.

## References

1. Takyar, S., R.P. Hickerson, and H.F. Noller, mRNA helicase activity of the ribosome. Cell, 2005. 120(1): p. 49–58.

2. Spinks, R.R., et al., DnaB helicase dynamics in bacterial DNA replication resolved by single-molecule studies. Nucleic Acids Research, 2021. 49(12): p. 6804–6816.

3. Chang, K.C. and J.D. Wen, Programmed −1 ribosomal frameshifting from the perspective of the conformational dynamics of mRNA and ribosomes. Comput Struct Biotechnol J, 2021. 19: p. 3580–3588.

4. Chen, J., et al., Dynamic pathways of −1 translational frameshifting. Nature, 2014. 512(7514): p. 328–32.

5. Mazauric, M.H., et al., Interaction of the HIV-1 frameshift signal with the ribosome. Nucleic Acids Res, 2009. 37(22): p. 7654–64.

6. Kim, H.K. and I. Tinoco, Jr., EF-G catalyzed translocation dynamics in the presence of ribosomal frameshifting stimulatory signals. Nucleic Acids Res, 2017. 45(5): p. 2865–2874.

7. Napthine, S., et al., Protein-directed ribosomal frameshifting temporally regulates gene expression. Nature Communications, 2017. 8(1): p. 15582.

8. Volkenborn, K., et al., The length of ribosomal binding site spacer sequence controls the production yield for intracellular and secreted proteins by Bacillus subtilis. Microbial Cell Factories, 2020. 19(1): p. 154.

9. Lin, Z., R.J. Gilbert, and I. Brierley, Spacer-length dependence of programmed −1 or −2 ribosomal frameshifting on a U6A heptamer supports a role for messenger RNA (mRNA) tension in frameshifting. Nucleic Acids Res, 2012. 40(17): p. 8674–89.

10. Chen, Y.-T., et al., Coordination among tertiary base pairs results in an efficient frameshift-stimulating RNA pseudoknot. Nucleic Acids Research, 2017. 45(10): p. 6011–6022.

11. Tholstrup, J., L.B. Oddershede, and M.A. Sørensen, mRNA pseudoknot structures can act as ribosomal roadblocks. Nucleic Acids Res, 2012. 40(1): p. 303–13.

12. Zhong, Z., et al., Mechanical unfolding kinetics of the SRV-1 gag-pro mRNA pseudoknot: possible implications for −1 ribosomal frameshifting stimulation. Scientific Reports, 2016. 6(1): p. 39549.

13. Nixon, P.L. and D.P. Giedroc, Equilibrium unfolding (folding) pathway of a model H-type pseudoknotted RNA: the role of magnesium ions in stability. Biochemistry, 1998. 37(46): p. 16116–29.

14. Qiu, H., et al., Thermodynamics of Folding of the RNA Pseudoknot of the T4 Gene 32 Autoregulatory Messenger RNA. Biochemistry, 1996. 35(13): p. 4176–4186.

15. Theimer, C.A. and D.P. Giedroc, Equilibrium unfolding pathway of an H-type RNA pseudoknot which promotes programmed −1 ribosomal frameshifting. J Mol Biol, 1999. 289(5): p. 1283–99.

16. Nixon, P.L., C.A. Theimer, and D.P. Giedroc, Thermodynamics of stabilization of RNA pseudoknots by cobalt(III) hexaammine. Biopolymers, 1999. 50(4): p. 443–58.

17. Theimer, C.A., et al., Non-nearest neighbor effects on the thermodynamics of unfolding of a model mRNA pseudoknot. J Mol Biol, 1998. 279(3): p. 545–64.

18. Wyatt, J.R., J.D. Puglisi, and I. Tinoco, Jr., RNA pseudoknots. Stability and loop size requirements. J Mol Biol, 1990. 214(2): p. 455–70.

19. Pallan, P.S., et al., Crystal structure of a luteoviral RNA pseudoknot and model for a minimal ribosomal frameshifting motif. Biochemistry, 2005. 44(34): p. 11315–22.

20. Su, L., et al., Minor groove RNA triplex in the crystal structure of a ribosomal frameshifting viral pseudoknot. Nature Structural Biology, 1999. 6(3): p. 285–92.

21. Jones, C.P. and A.R. Ferré-D’Amaré, Crystal structure of the severe acute respiratory syndrome coronavirus 2 (SARS-CoV-2) frameshifting pseudoknot. Rna, 2022. 28(2): p. 239–249.

22. Frank, J., et al., A model of protein synthesis based on cryo-electron microscopy of the E. coli ribosome. Nature, 1995. 376(6539): p. 441–4.

23. Cornish, P.V., M. Hennig, and D.P. Giedroc, A loop 2 cytidine-stem 1 minor groove interaction as a positive determinant for pseudoknot-stimulated −1 ribosomal frameshifting. Proc Natl Acad Sci USA, 2005. 102(36): p. 12694–9.

24. Cornish, P.V., S.N. Stammler, and D.P. Giedroc, The global structures of a wild-type and poorly functional plant luteoviral mRNA pseudoknot are essentially identical. RNA, 2006. 12(11): p. 1959–69.

25. Holland, J.A., et al., An examination of coaxial stacking of helical stems in a pseudoknot motif: the gene 32 messenger RNA pseudoknot of bacteriophage T2. Rna, 1999. 5(2): p. 257–71.

26. Shen, L.X. and I. Tinoco, The structure of an RNA pseudoknot that causes efficient frameshifting in mouse mammary tumor virus. J Mol Biol, 1995. 247(5): p. 963–78.

27. Puglisi, J.D., J.R. Wyatt, and I. Tinoco, Jr., Conformation of an RNA pseudoknot. J Mol Biol, 1990. 214(2): p. 437–53.

28. Green, L., et al., Characterization of the mechanical unfolding of RNA pseudoknots. J Mol Biol, 2008. 375(2): p. 511–28.

29. Hansen, T.M., et al., Correlation between mechanical strength of messenger RNA pseudoknots and ribosomal frameshifting. Proc Natl Acad Sci USA, 2007. 104(14): p. 5830–5.

30. Chen, G., et al., Triplex structures in an RNA pseudoknot enhance mechanical stability and increase efficiency of −1 ribosomal frameshifting. Proc Natl Acad Sci U S A, 2009. 106(31): p. 12706–11.

31. Neupane, K., et al., Structural dynamics of single SARS-CoV-2 pseudoknot molecules reveal topologically distinct conformers. Nat Commun, 2021. 12(1): p. 4749.

32. Ritchie, D.B., D.A. Foster, and M.T. Woodside, Programmed −1 frameshifting efficiency correlates with RNA pseudoknot conformational plasticity, not resistance to mechanical unfolding. Proc Natl Acad Sci U S A, 2012. 109(40): p. 16167–72.

33. Zhang, X., et al., Mimicking Ribosomal Unfolding of RNA Pseudoknot in a Protein Channel. J Am Chem Soc, 2015. 137(50): p. 15742–52.

34. Napthine, S., et al., The role of RNA pseudoknot stem 1 length in the promotion of efficient −1 ribosomal frameshifting 11Edited by D. E. Draper. Journal of Molecular Biology, 1999. 288(3): p. 305–320.

35. Kim, Y.G., et al., Specific mutations in a viral RNA pseudoknot drastically change ribosomal frameshifting efficiency. Proc Natl Acad Sci U S A, 1999. 96(25): p. 14234–9.

36. Desai, V.P., et al., Co-temporal Force and Fluorescence Measurements Reveal a Ribosomal Gear Shift Mechanism of Translation Regulation by Structured mRNAs. Molecular Cell, 2019. 75(5): p. 1007–1019.e5.

37. Qu, X., et al., The ribosome uses two active mechanisms to unwind messenger RNA during translation. Nature, 2011. 475(7354): p. 118–21.

38. Qu, X., et al., Ribosomal protein S1 unwinds double-stranded RNA in multiple steps. Proc Natl Acad Sci U S A, 2012. 109(36): p. 14458–63.

39. Chen, C., et al., Translation by Single Ribosomes Through mRNA Secondary Structures. Biophysical Journal, 2012. 102(3): p. 67a–68a.

40. Namy, O., et al., A mechanical explanation of RNA pseudoknot function in programmed ribosomal frameshifting. Nature, 2006. 441(7090): p. 244–7.

41. Lund, P.E., et al., Protein unties the pseudoknot: S1-mediated unfolding of RNA higher order structure. Nucleic Acids Research, 2019. 48(4): p. 2107–2125.

42. Mazauric, M.H., et al., Footprinting analysis of BWYV pseudoknot-ribosome complexes. RNA, 2009. 15(9): p. 1775–86.

43. Thulson, E., et al., An RNA pseudoknot stimulates HTLV-1 pro-pol programmed - 1 ribosomal frameshifting. Rna, 2020. 26(4): p. 512–528.

44. Krokhotin, A., et al., Direct identification of base-paired RNA nucleotides by correlated chemical probing. Rna, 2017. 23(1): p. 6–13.

45. Mitchell, D., III, et al., Mutation signature filtering enables high-fidelity RNA structure probing at all four nucleobases with DMS. Nucleic Acids Research, 2023. 51(16): p. 8744–8757.

46. Lawley, P.D. and P. Brookes, FURTHER STUDIES ON THE ALKYLATION OF NUCLEIC ACIDS AND THEIR CONSTITUENT NUCLEOTIDES. Biochemical Journal, 1963. 89(1): p. 127–138.

47. Mustoe, A.M., et al., RNA base-pairing complexity in living cells visualized by correlated chemical probing. Proceedings of the National Academy of Sciences, 2019. 116(49): p. 24574–24582.

48. Zubradt, M., et al., DMS-MaPseq for genome-wide or targeted RNA structure probing in vivo. Nature Methods, 2017. 14(1): p. 75–82.

49. Tomezsko, P., H. Swaminathan, and S. Rouskin, Viral RNA structure analysis using DMS-MaPseq. Methods, 2020. 183: p. 68–75.

50. Sanduni Deenalattha, D.H., et al., Characterizing 3D RNA structural features from DMS reactivity. bioRxiv, 2025: p. 2024.11.21.624766.

51. Saha, K. and G. Ghosh, Chemical Probing of RNA Structure In Vivo Using SHAPE-MaP and DMS-MaP, in RNA-Protein Complexes and Interactions: Methods and Protocols, R.-J. Lin, Editor. 2023, Springer US: New York, NY. p. 81–93.

52. Cornish, P.V., et al., Spontaneous intersubunit rotation in single ribosomes. Mol Cell, 2008. 30(5): p. 578–88.

53. Forino, N.M., et al., Telomerase RNA structural heterogeneity in living human cells detected by DMS-MaPseq. Nature Communications, 2025. 16(1): p. 925.

54. Jenner, L.B., et al., Structural aspects of messenger RNA reading frame maintenance by the ribosome. Nat Struct Mol Biol, 2010. 17(5): p. 555–60.

55. Qin, P., et al., Structured mRNA induces the ribosome into a hyper-rotated state. EMBO Rep, 2014. 15(2): p. 185–90.

56. Shebl, B., et al., The influence of downstream structured elements within mRNA on the dynamics of intersubunit rotation in ribosomes. bioRxiv, 2024: p. 2024.10.17.618931.

57. Ritchey, L.E., et al., Structure-seq2: sensitive and accurate genome-wide profiling of RNA structure in vivo. Nucleic Acids Research, 2017. 45(14): p. e135–e135.

58. Choudhary, K., et al., dStruct: identifying differentially reactive regions from RNA structurome profiling data. Genome Biology, 2019. 20(1): p. 40.

59. Kwok, C.K., et al., The RNA structurome: transcriptome-wide structure probing with next-generation sequencing. Trends in Biochemical Sciences, 2015. 40(4): p. 221–232.

60. Kuksa, P.P., et al., HiPR: High-throughput probabilistic RNA structure inference. Computational and Structural Biotechnology Journal, 2020. 18: p. 1539–1547.

61. Incarnato, D., et al., RNA Framework: an all-in-one toolkit for the analysis of RNA structures and post-transcriptional modifications. Nucleic Acids Research, 2018. 46(16): p. e97–e97.

62. Atkins, J.F. and R.F. Gesteland, Intricacies of ribosomal frameshifting. Nat Struct Biol, 1999. 6(3): p. 206–7.

63. Walter, A.E., et al., Coaxial stacking of helixes enhances binding of oligoribonucleotides and improves predictions of RNA folding. Proceedings of the National Academy of Sciences, 1994. 91(20): p. 9218–9222.

64. Walter, A.E. and D.H. Turner, Sequence dependence of stability for coaxial stacking of RNA helixes with Watson-Crick base paired interfaces. Biochemistry, 1994. 33(42): p. 12715–12719.

65. Cornish, P.V. and D.P. Giedroc, Pairwise coupling analysis of helical junction hydrogen bonding interactions in luteoviral RNA pseudoknots. Biochemistry, 2006. 45(37): p. 11162–71.

66. Peattie, D.A. and W. Gilbert, Chemical probes for higher-order structure in RNA. Proceedings of the National Academy of Sciences, 1980. 77(8): p. 4679–4682.

67. Plant, E.P. and J.D. Dinman, Torsional restraint: a new twist on frameshifting pseudoknots. Nucleic Acids Res, 2005. 33(6): p. 1825–33.

68. Giedroc, D.P., P.V. Cornish, and M. Hennig, Detection of scalar couplings involving 2’-hydroxyl protons across hydrogen bonds in a frameshifting mRNA pseudoknot. J Am Chem Soc, 2003. 125(16): p. 4676–7.

69. Jones, C.P. and A.R. Ferré-D’Amaré, Structural switching dynamically controls the doubly pseudoknotted Rous sarcoma virus–programmed ribosomal frameshifting element. Proceedings of the National Academy of Sciences, 2025. 122(14): p. e2418418122.

70. Xie, P., Model of the pathway of −1 frameshifting: Long pausing. Biochemistry and Biophysics Reports, 2016. 5: p. 408–424.

71. Kim, H.K., et al., A frameshifting stimulatory stem loop destabilizes the hybrid state and impedes ribosomal translocation. Proc Natl Acad Sci U S A, 2014. 111(15): p. 5538–43.

72. Choi, J., et al., The energy landscape of −1 ribosomal frameshifting. Science Advances. 6(1): p. eaax6969.

73. Adamski, F.M., B.C. Donly, and W.P. Tate, Competition between frameshifting, termination and suppression at the frameshift site in the Escherichia coli release factor-2 mRNA. Nucleic Acids Res, 1993. 21(22): p. 5074–8.

74. Bao, C., et al., Specific length and structure rather than high thermodynamic stability enable regulatory mRNA stem-loops to pause translation. Nature Communications, 2022. 13(1): p. 988.

75. Wen, J.D., et al., Following translation by single ribosomes one codon at a time. Nature, 2008. 452(7187): p. 598–603.

76. Ritchie, D.B., D.A.N. Foster, and M.T. Woodside, Programmed −1 frameshifting efficiency correlates with RNA pseudoknot conformational plasticity, not resistance to mechanical unfolding. Proceedings of the National Academy of Sciences, 2012. 109(40): p. 16167–16172.

77. Hsu, C.-F., et al., Formation of frameshift-stimulating RNA pseudoknots is facilitated by remodeling of their folding intermediates. Nucleic Acids Research, 2021. 49(12): p. 6941–6957.

78. Petrov, A., et al., Chapter Seventeen - RNA Purification by Preparative Polyacrylamide Gel Electrophoresis, in Methods in Enzymology, J. Lorsch, Editor. 2013, Academic Press. p. 315–330.

79. Staehelin, T. and D.R. Maglott, [47] Preparation of Escherchia coli ribosomal subunits active in polypeptide synthesis, in Methods in Enzymology. 1971, Academic Press. p. 449–456.

80. Smola, M.J., et al., *Selective 2*′*-hydroxyl acylation analyzed by primer extension and mutational profiling (SHAPE-MaP) for direct, versatile and accurate RNA structure analysis*. Nature Protocols, 2015. 10(11): p. 1643–1669.

81. Leontis, N.B., J. Stombaugh, and E. Westhof, *The non*-*Watson–Crick base pairs and their associated isostericity matrices*. Nucleic Acids Research, 2002. 30(16): p. 3497–3531.

