## Supplemental Figure S1 and S2 for "Ribosome-induced mRNA pseudoknot interactions visualized by DMS MaP-Seq"

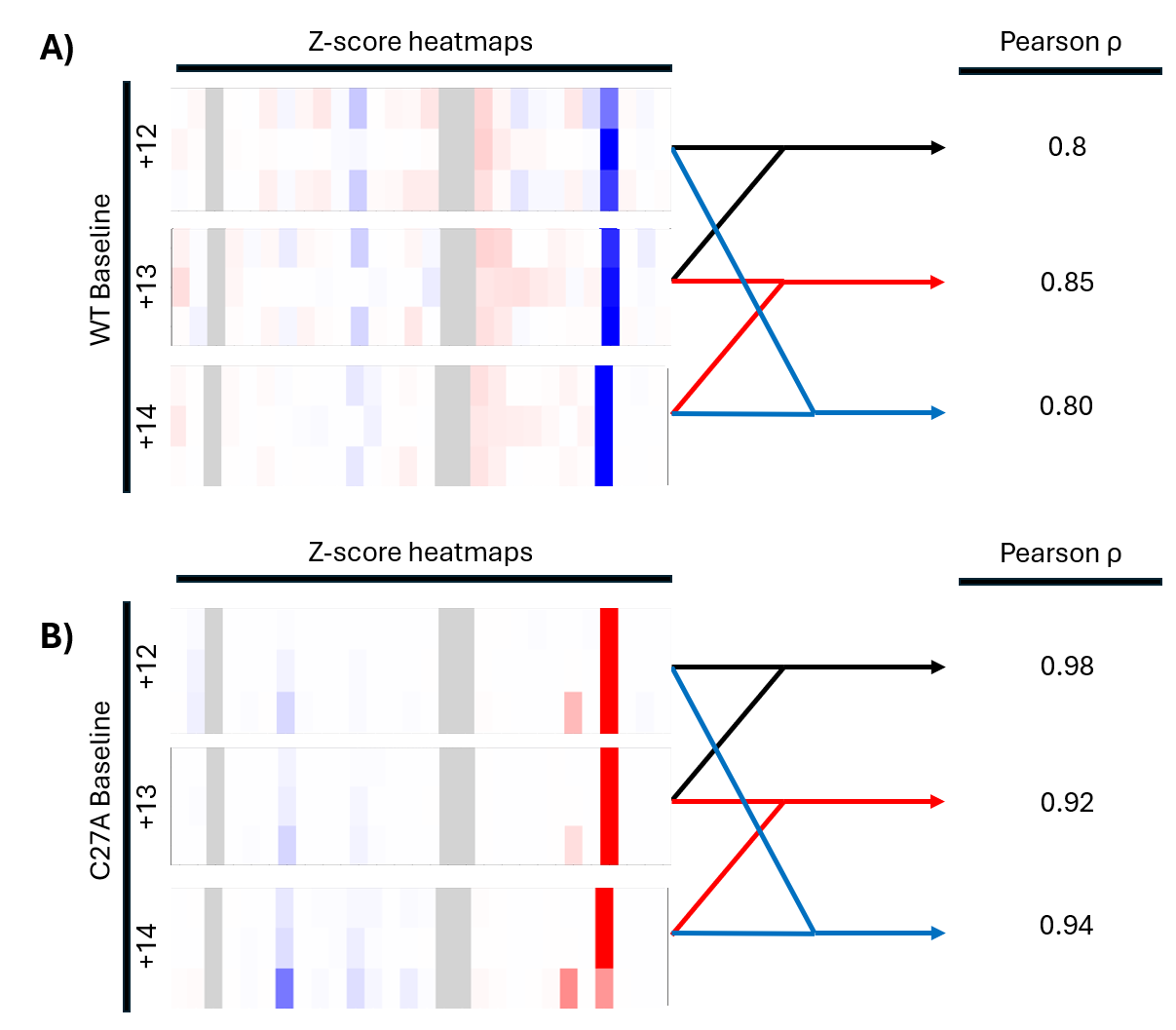

***Figure S1: Similarity between variable linker RNAs in reproducing experimental results.*** *A). Each heatmap contains z score differences as calculated from +12, +13, and +14. X axis is sequence of ScYLV. Y axis is experimental conditions (aP-tRNA, daP-tRNA, PRE) The heatmaps are then compared to one another to generate Pearson ρ.*

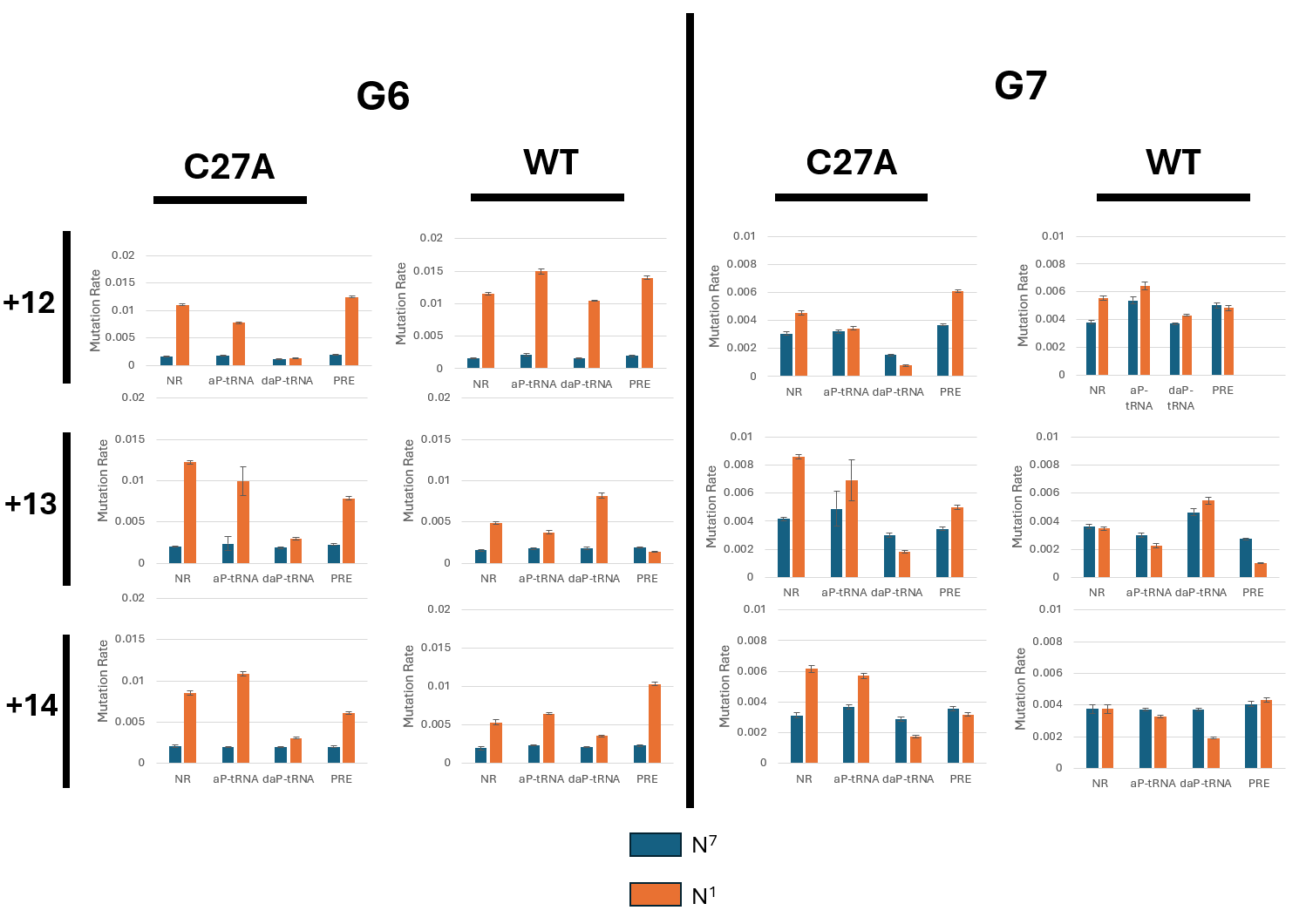

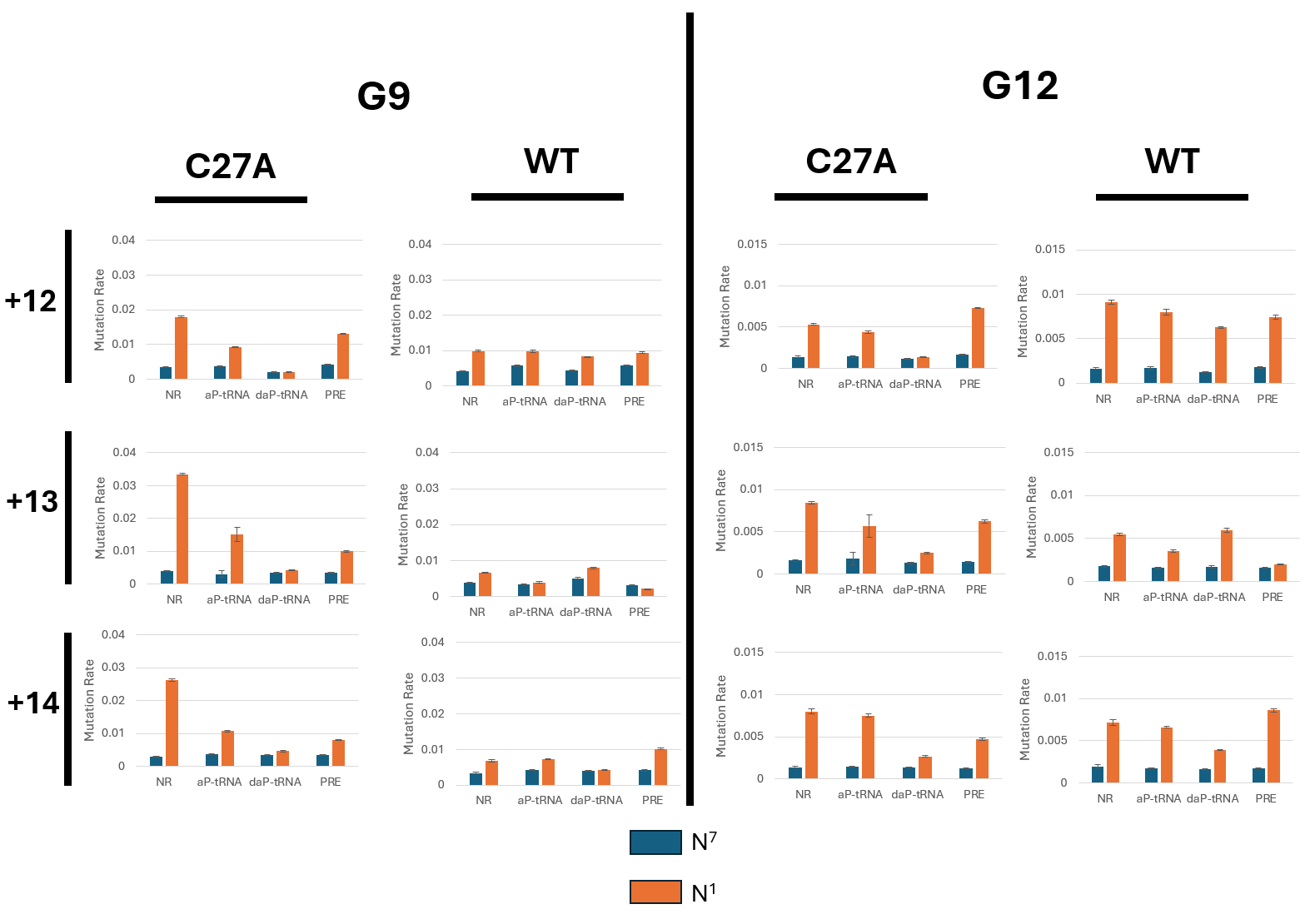

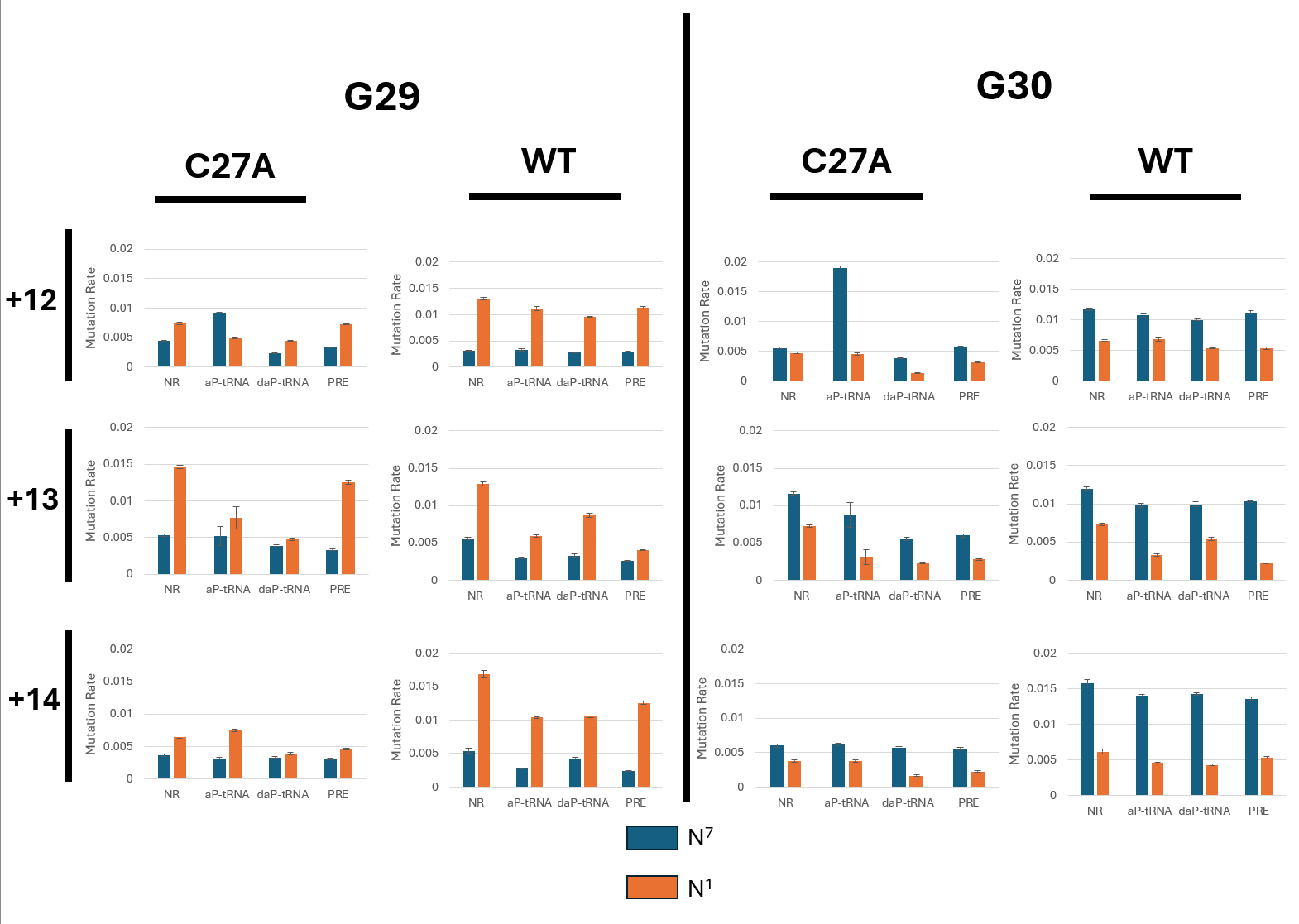

*Figure S2: Bar graphs of N^7^ and N^1^ modification rates of G nucleotides present in ScYLVPK across all ribosomal states [NR, aP-tRNA, daP-tRNA, PRE) , linker lengths (+12, +13, +14) , and RNAs (C27A and WT). N^7^ modification is depicted in blue and N^1^ modification rate is depicted in orange.*
