## Supplementary figures and images for "Ribosome-induced mRNA pseudoknot interactions visualized by DMS MaP-Seq"

### Target profiles 1

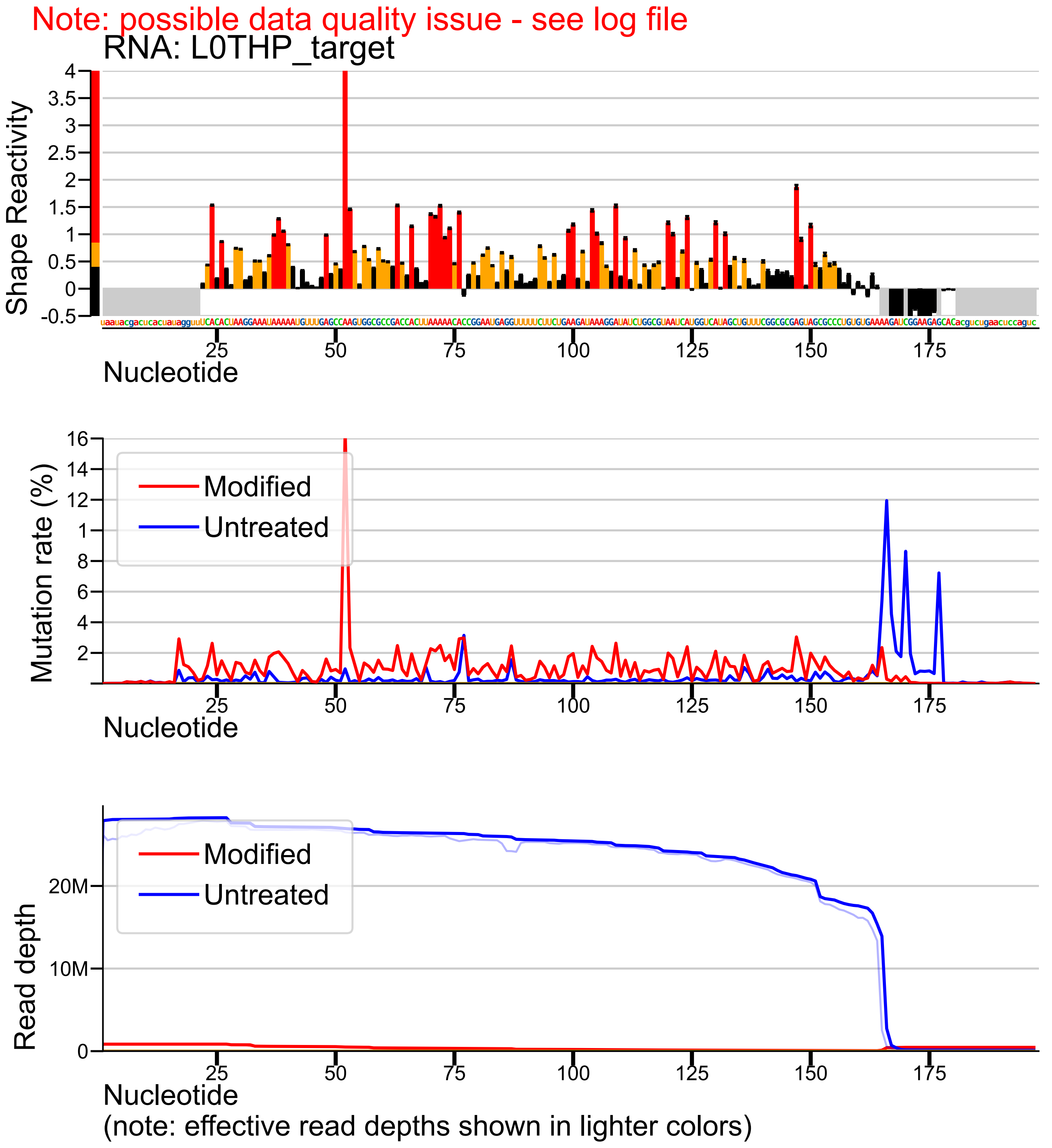

### Target profiles 2

Note: possible data quality issue - see log file

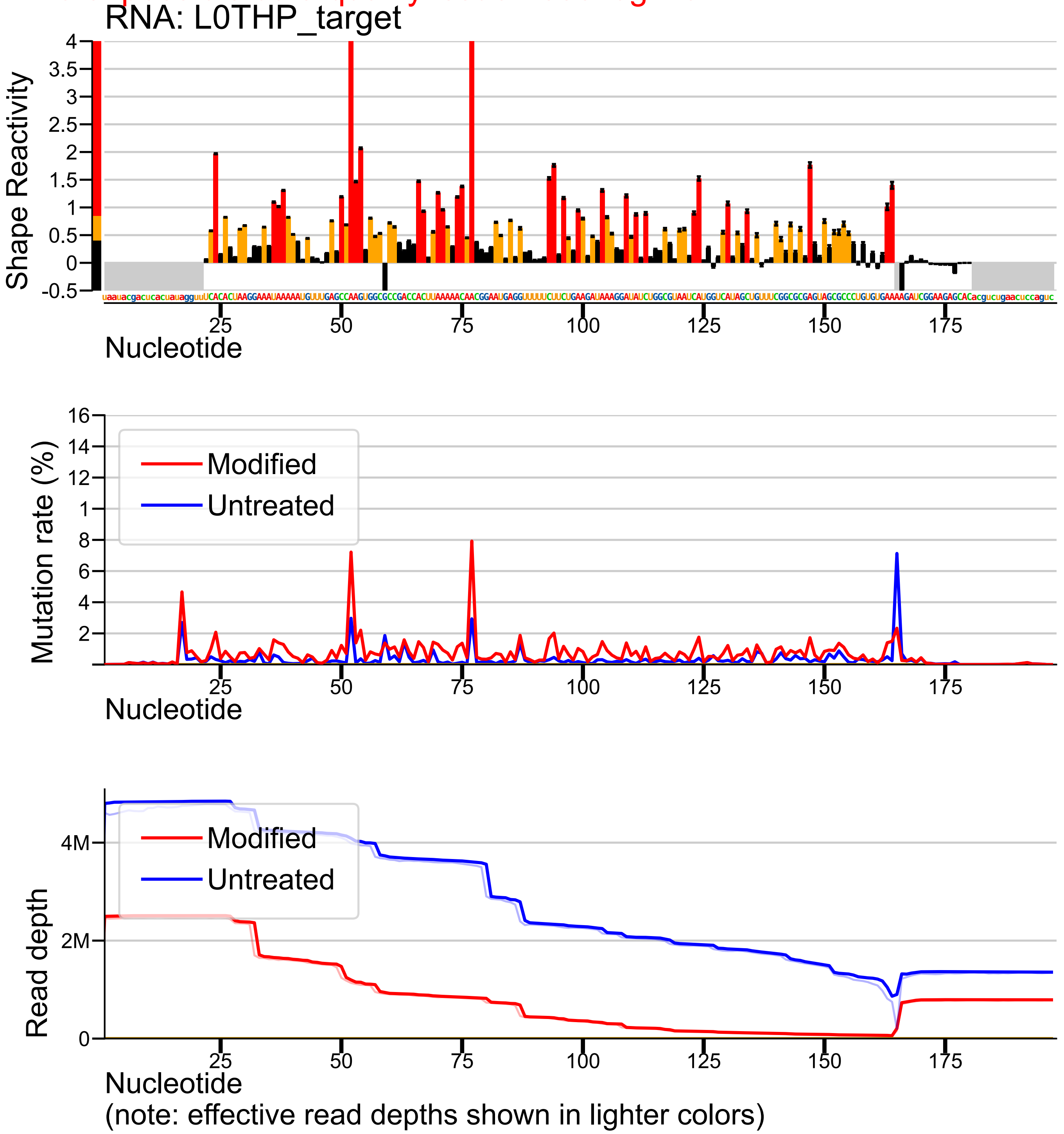

### Target profiles 3

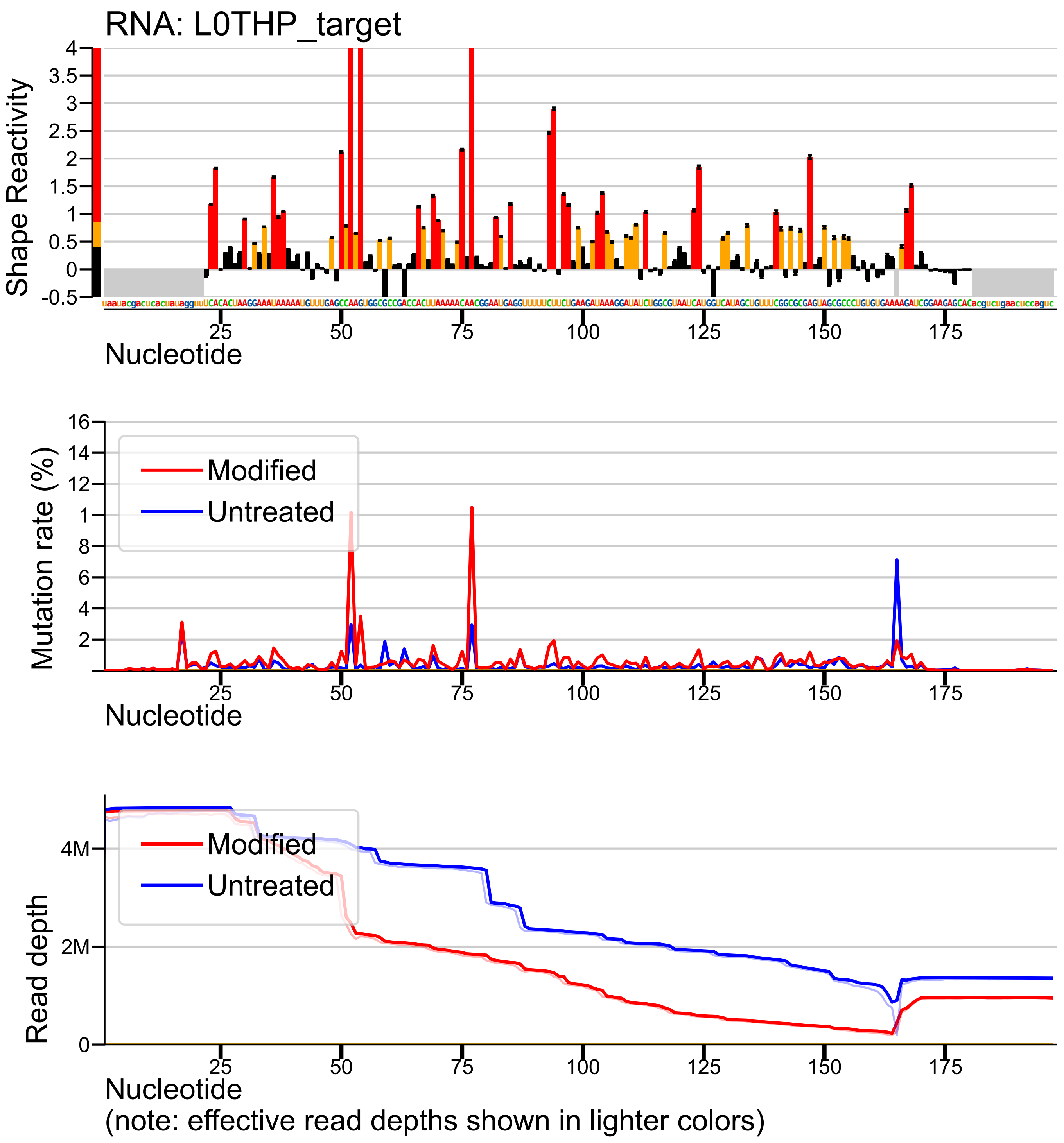

### Target profiles 4

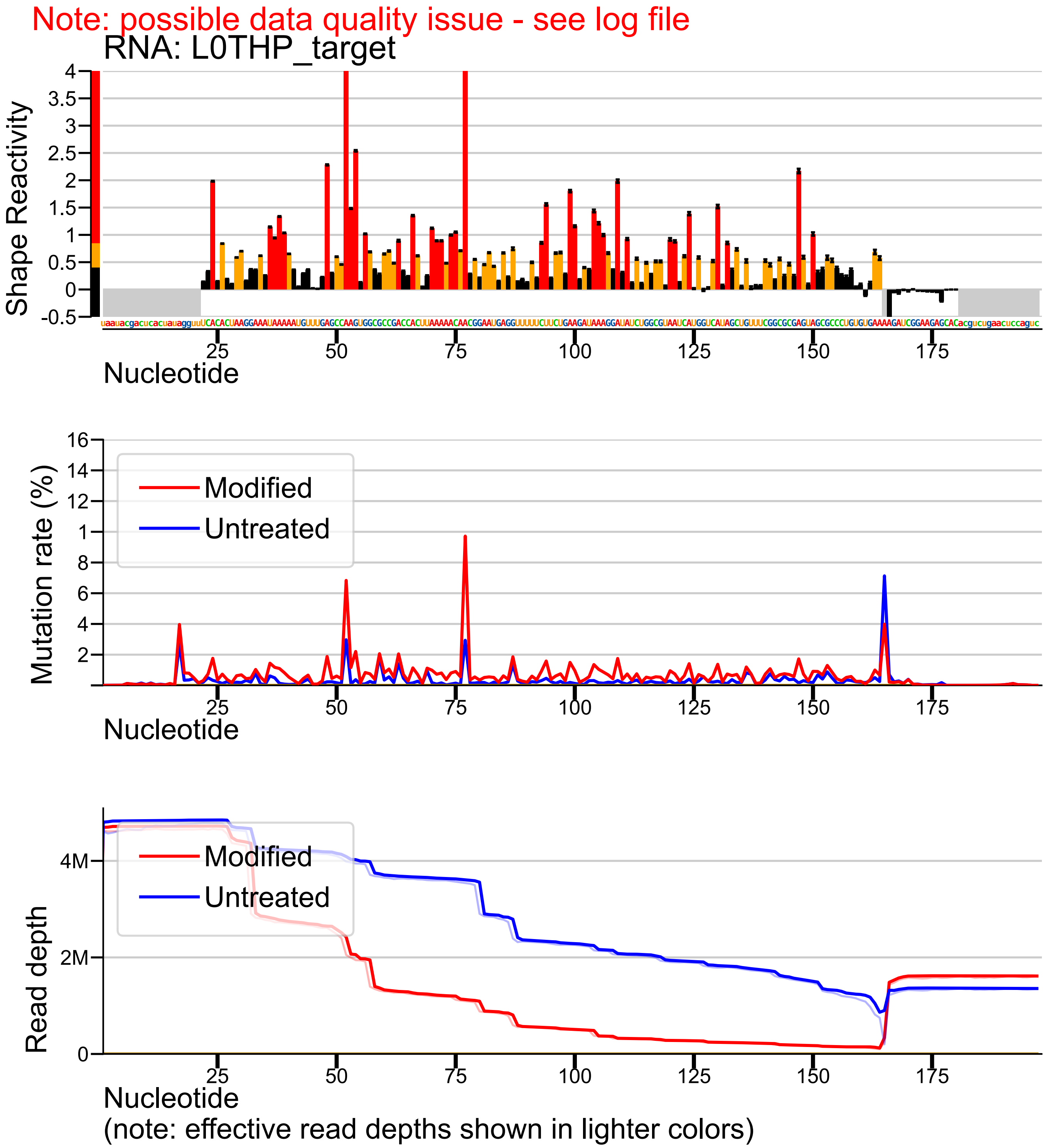

### Target profiles 5

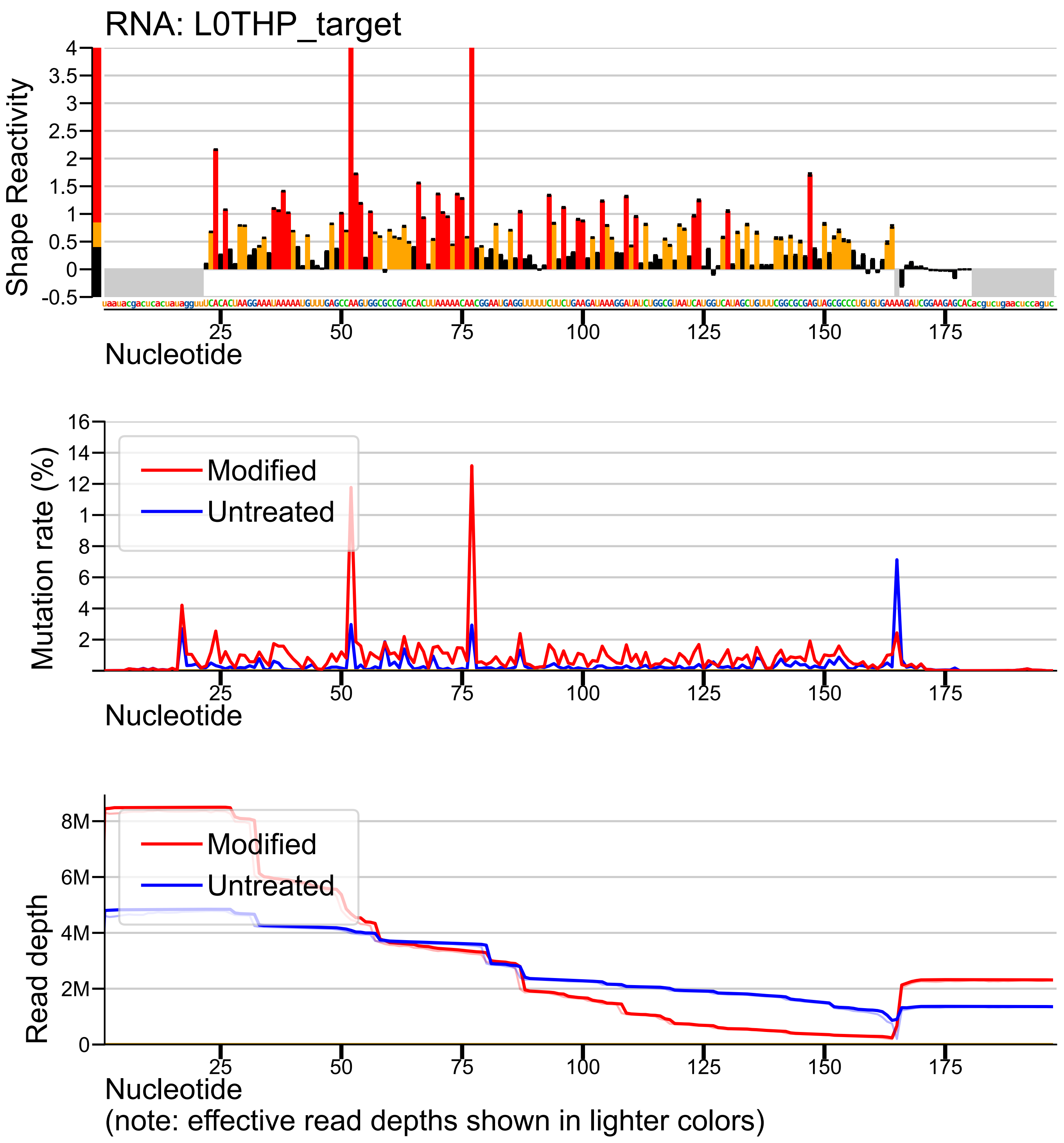

### Target profiles 6

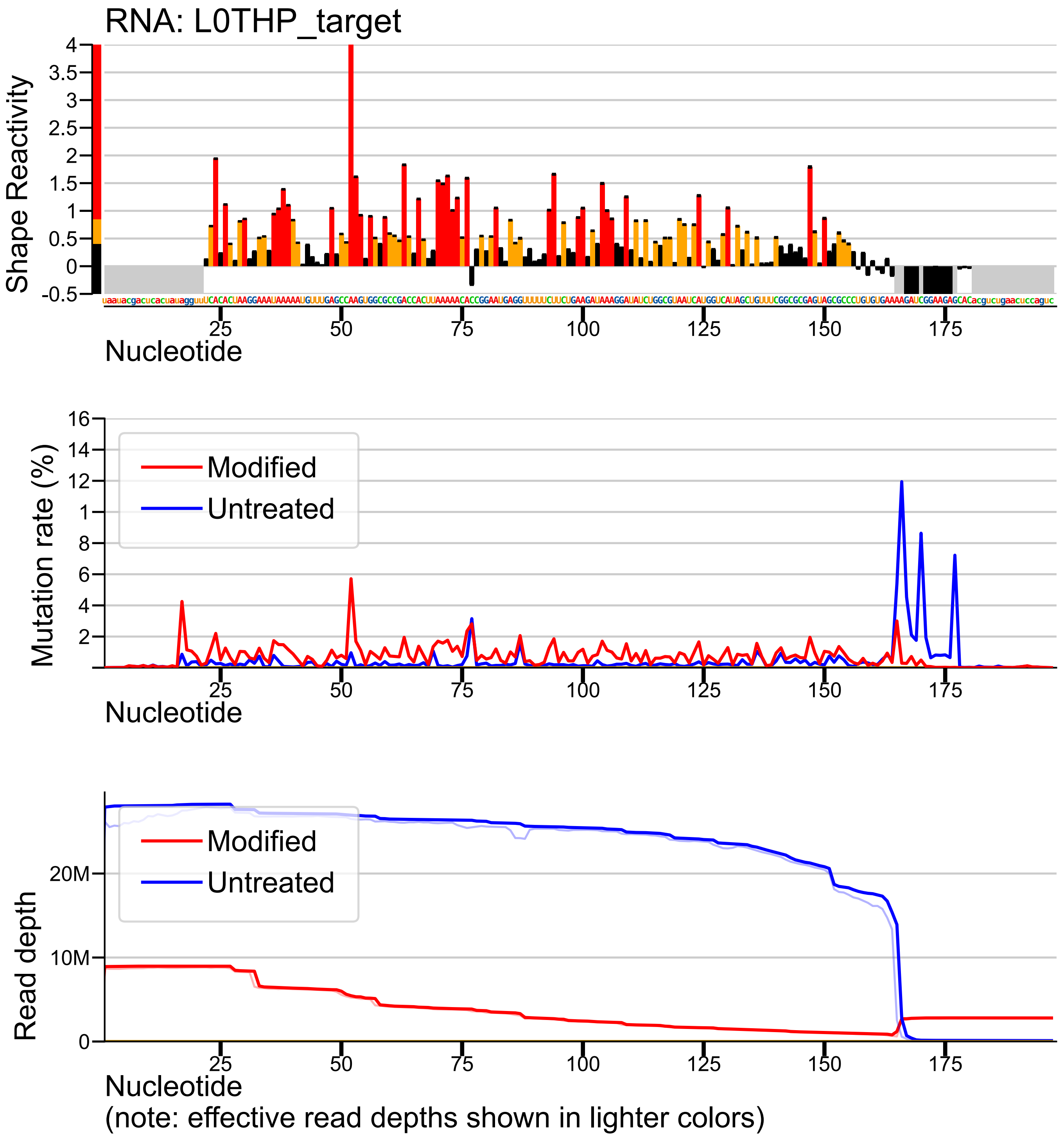

### Target profiles 7

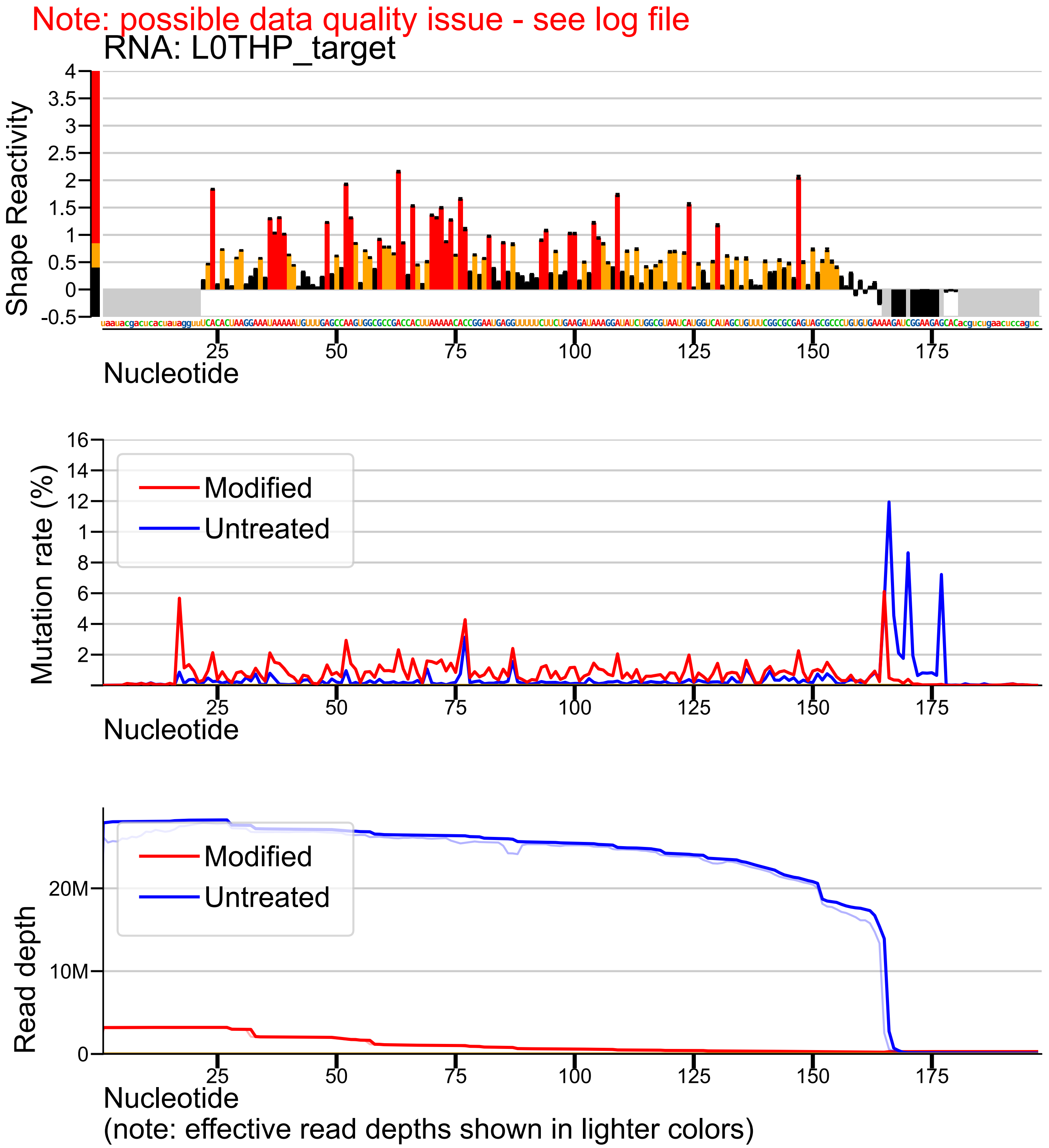

### Target profiles 9

Note: possible data quality issue - see log file

RNA: L1THP_target

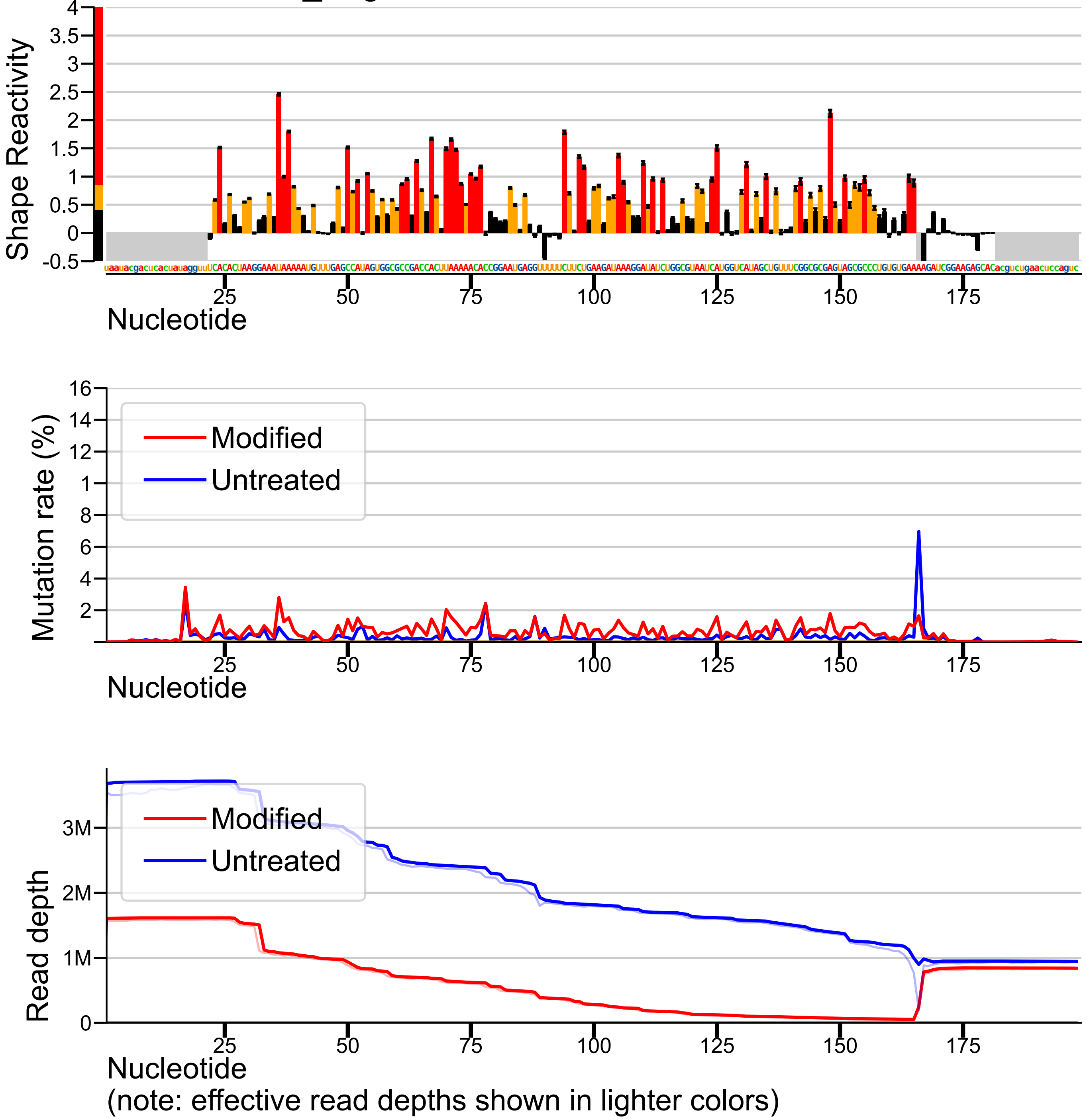

### Target profiles 10

Note: possible data quality issue - see log file

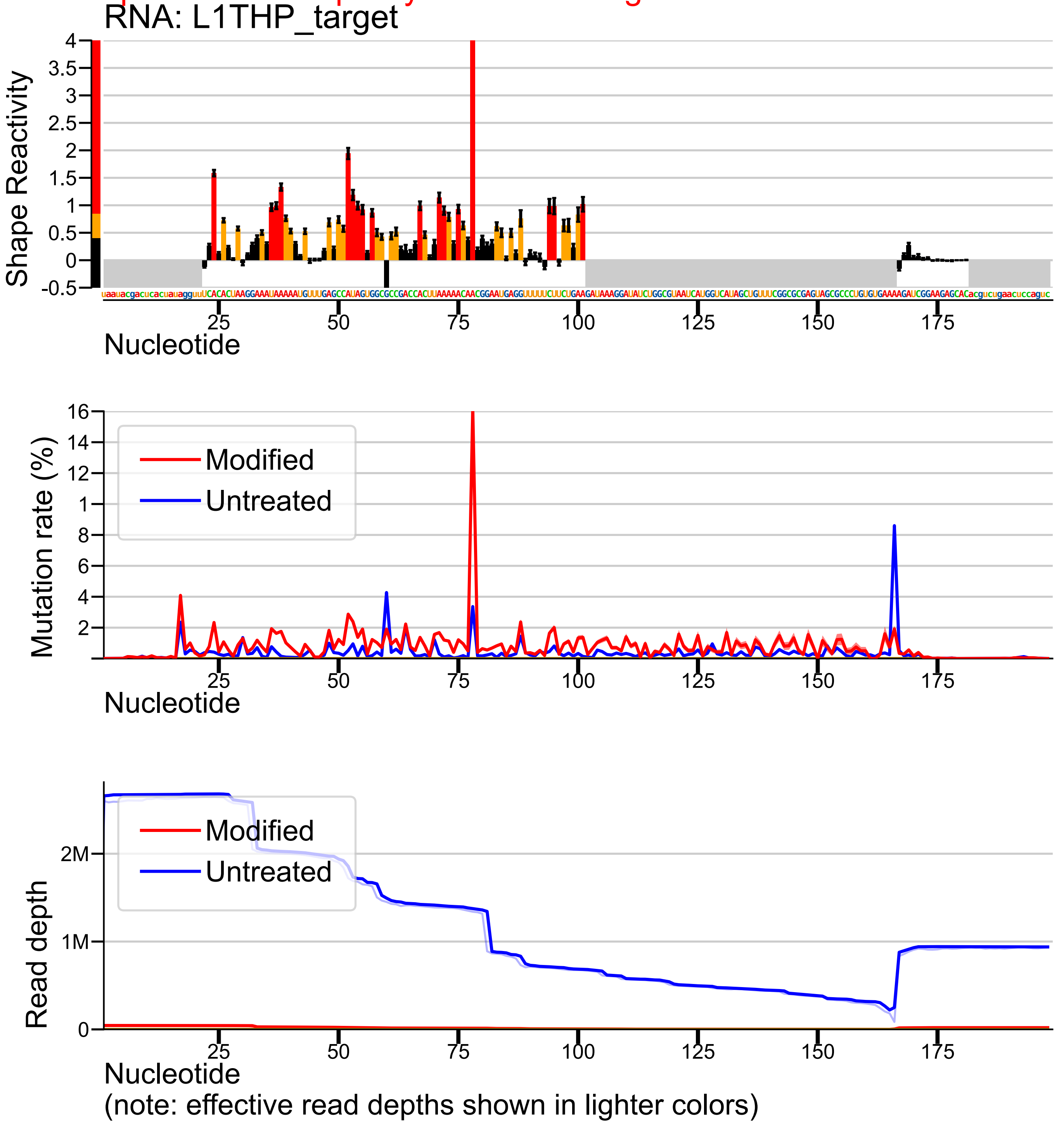

### Target profiles 11

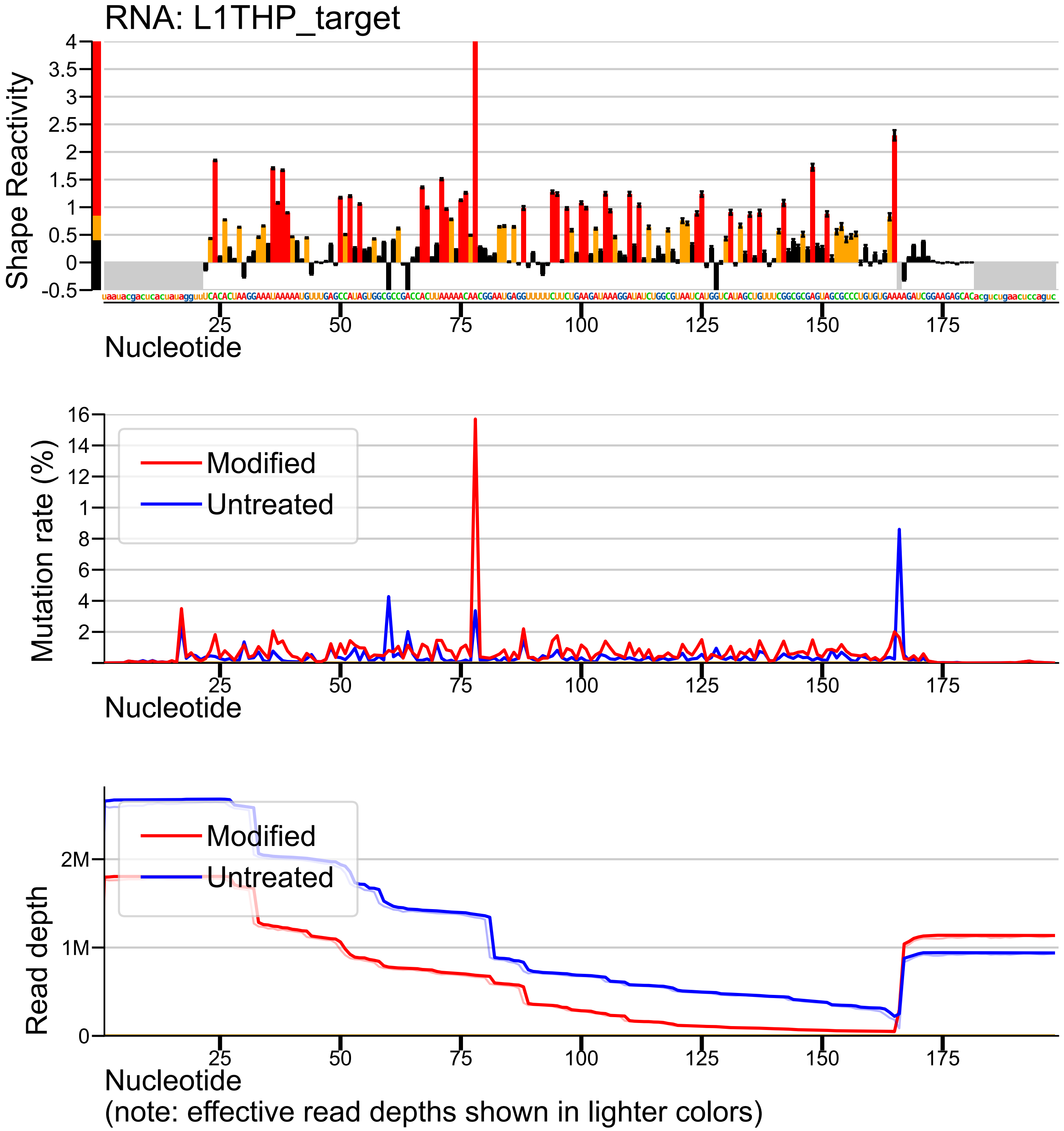

### Target profiles 13

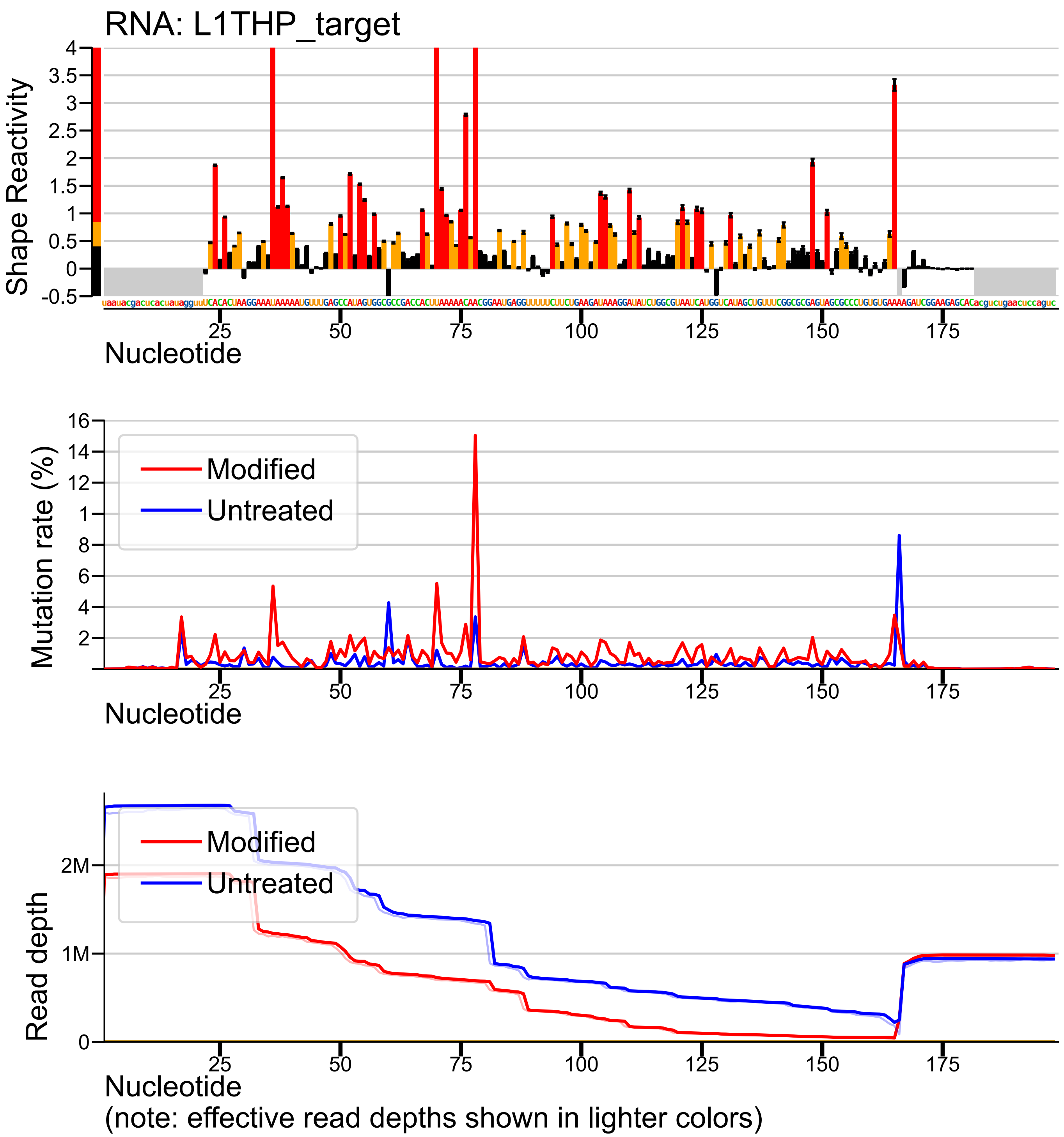

### Target profiles 14

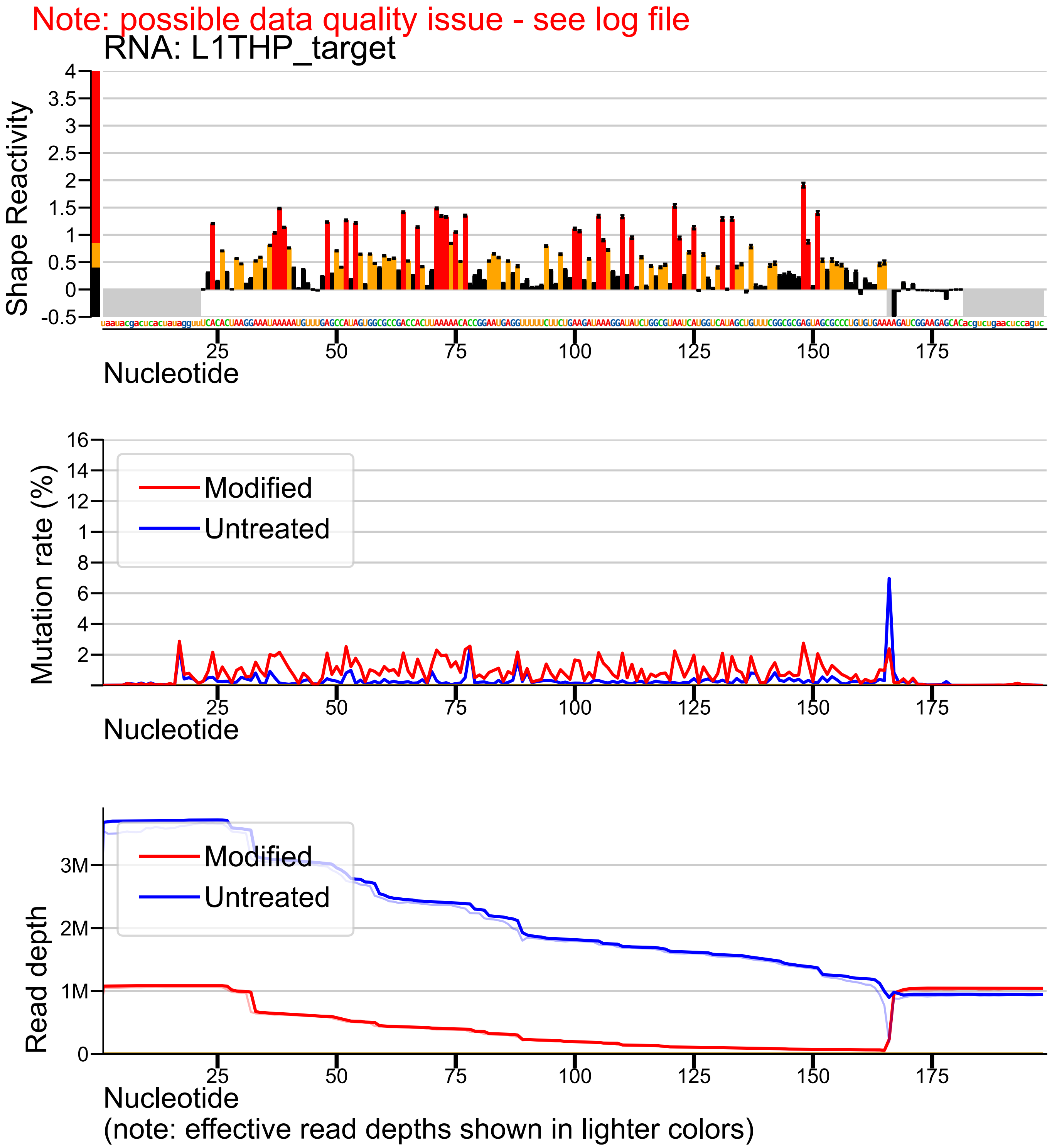

### Target profiles 15

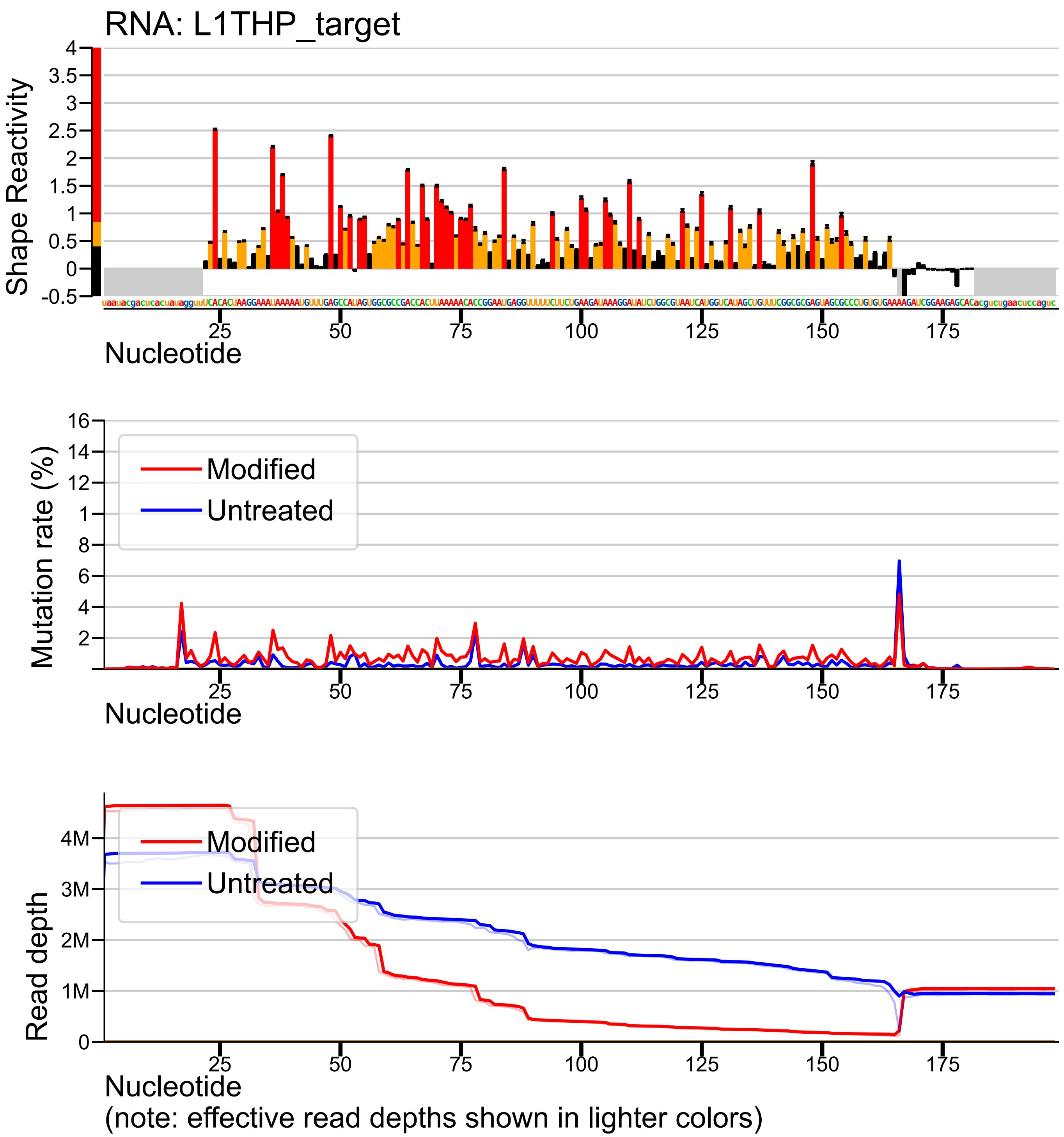

### Target profiles 16

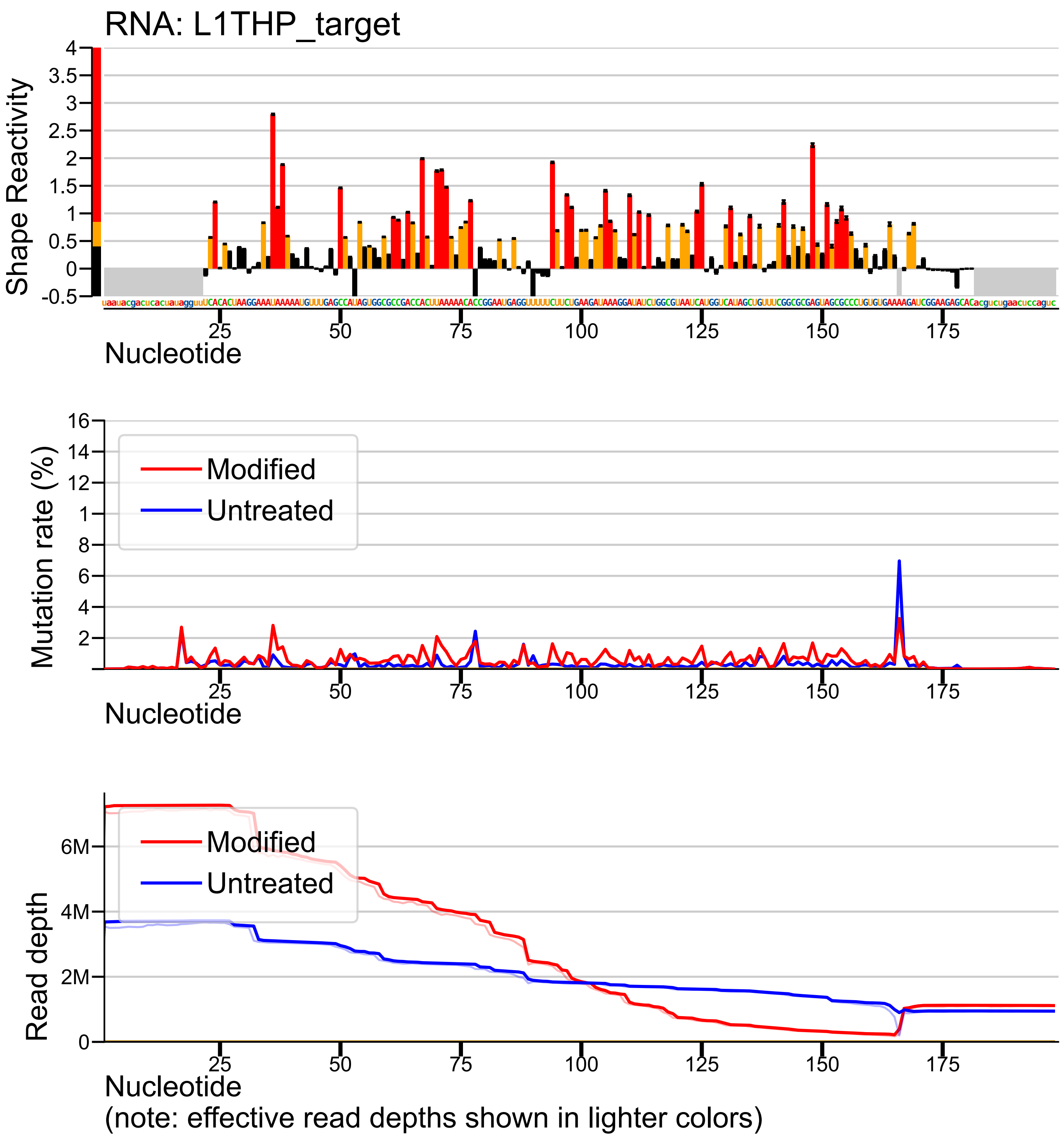

### Target profiles 18

Note: possible data quality issue - see log file

RNA: L2THP_target

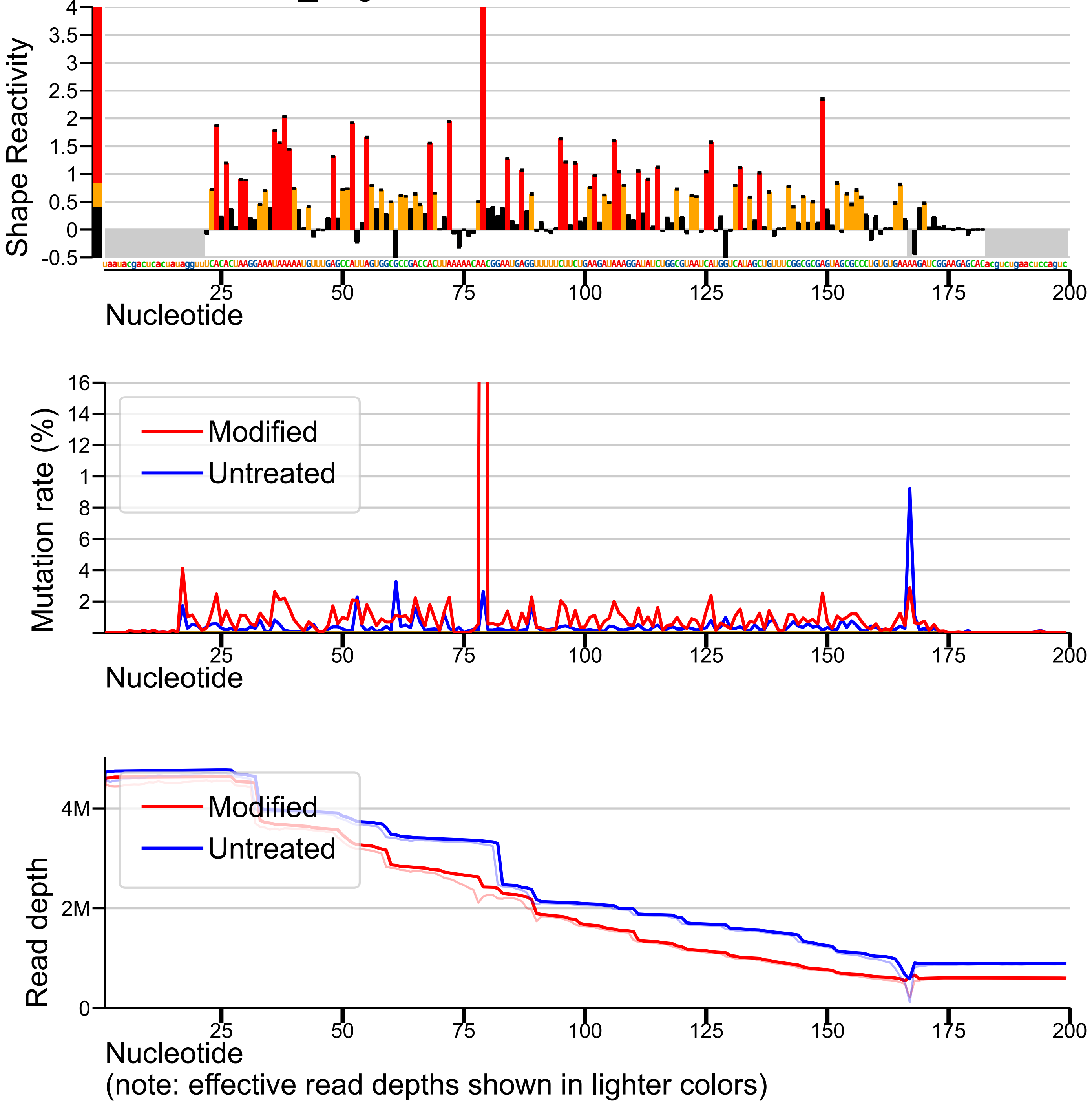

### Target profiles 19

Note: possible data quality issue - see log file

RNA: L2THP_target

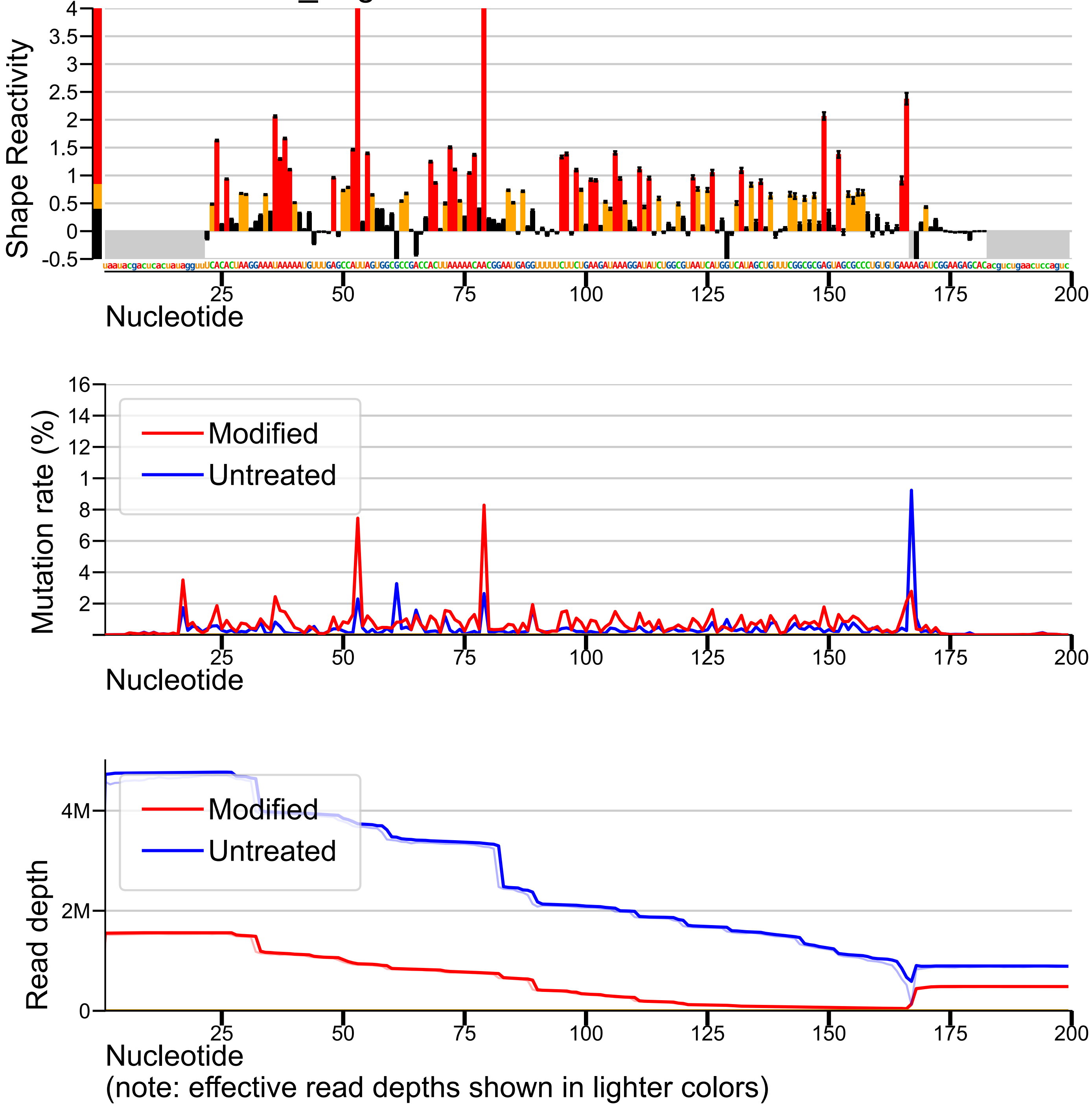

### Target profiles 20

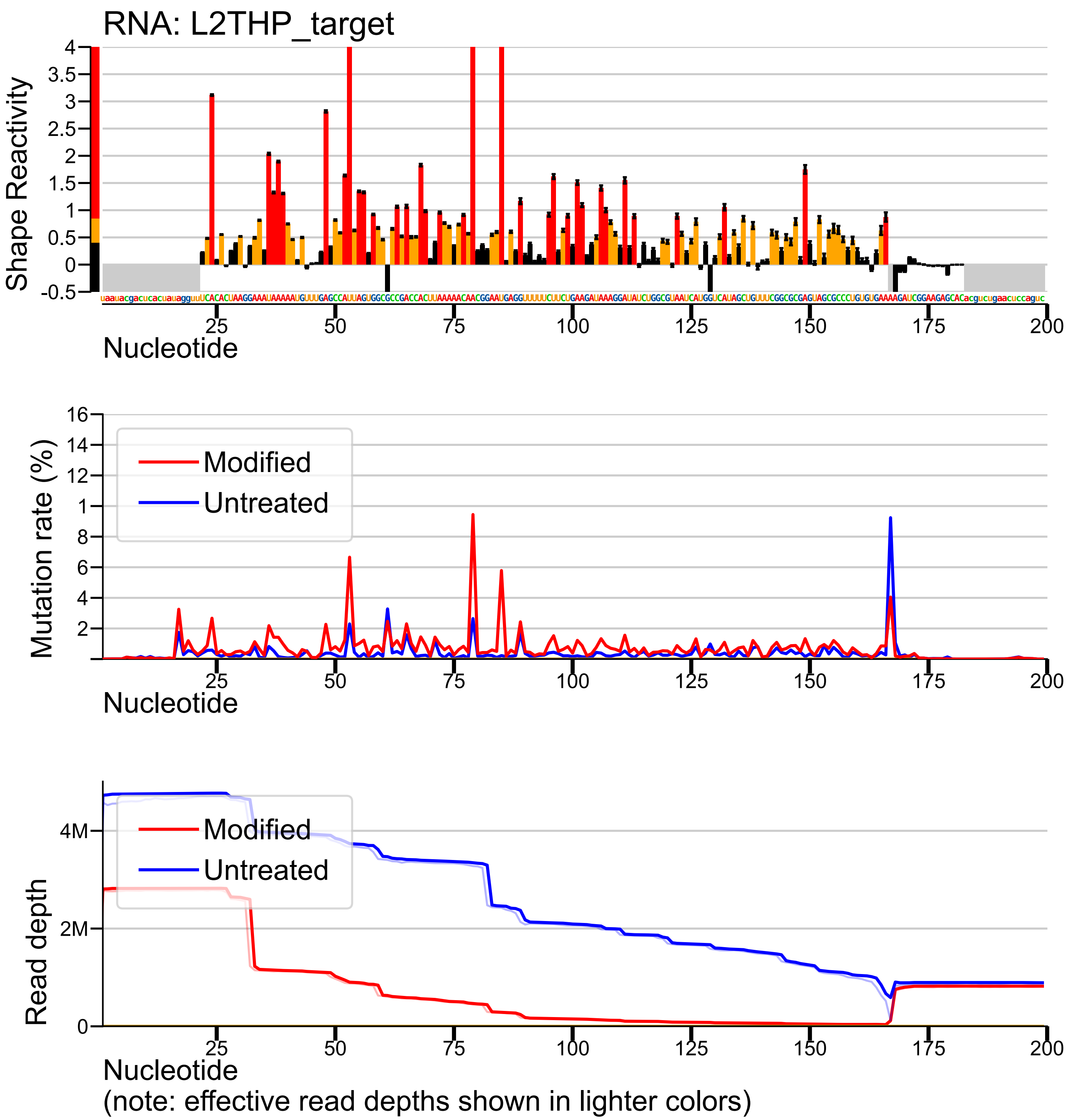

### Target profiles 22

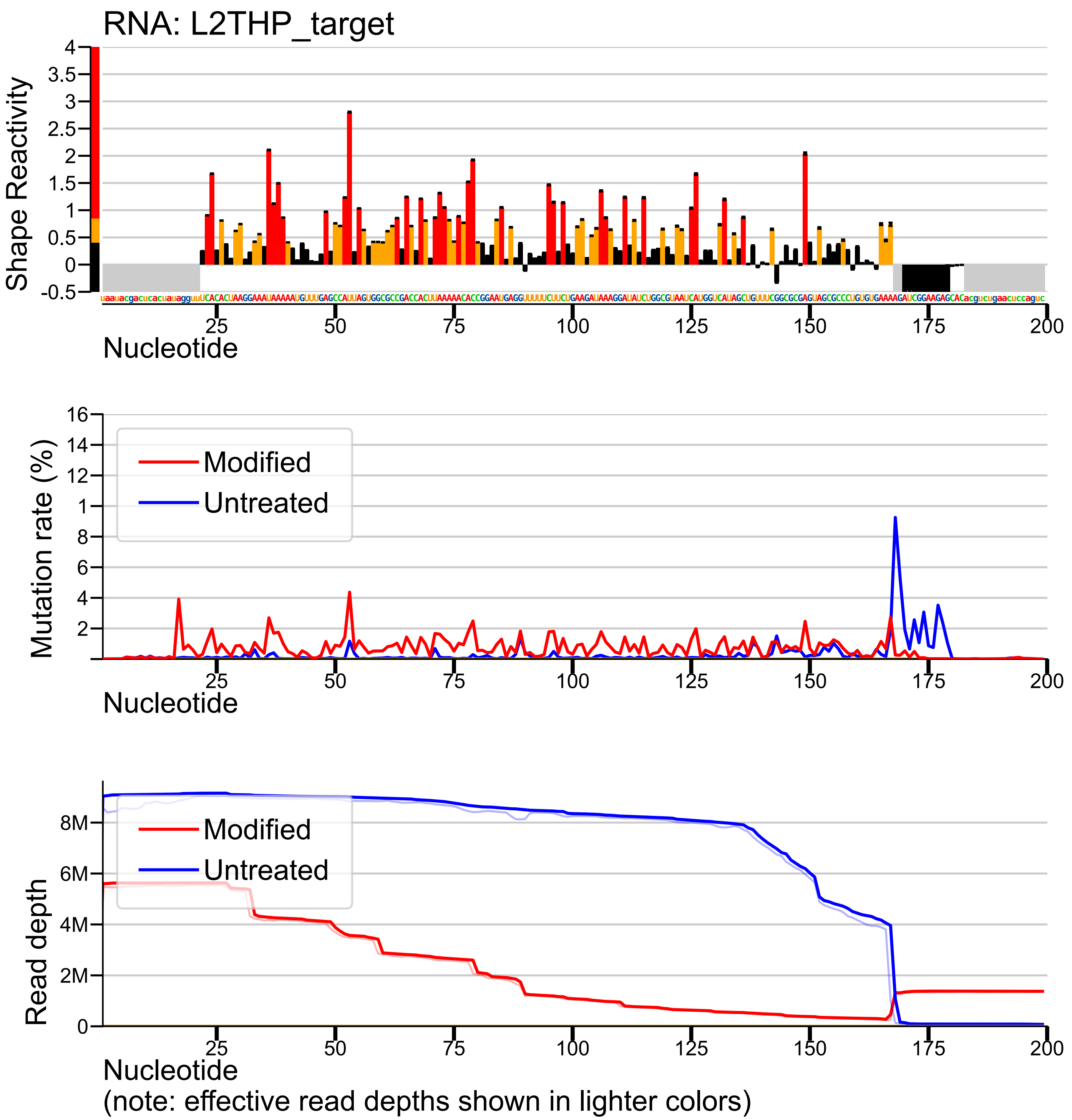

### Target profiles 23

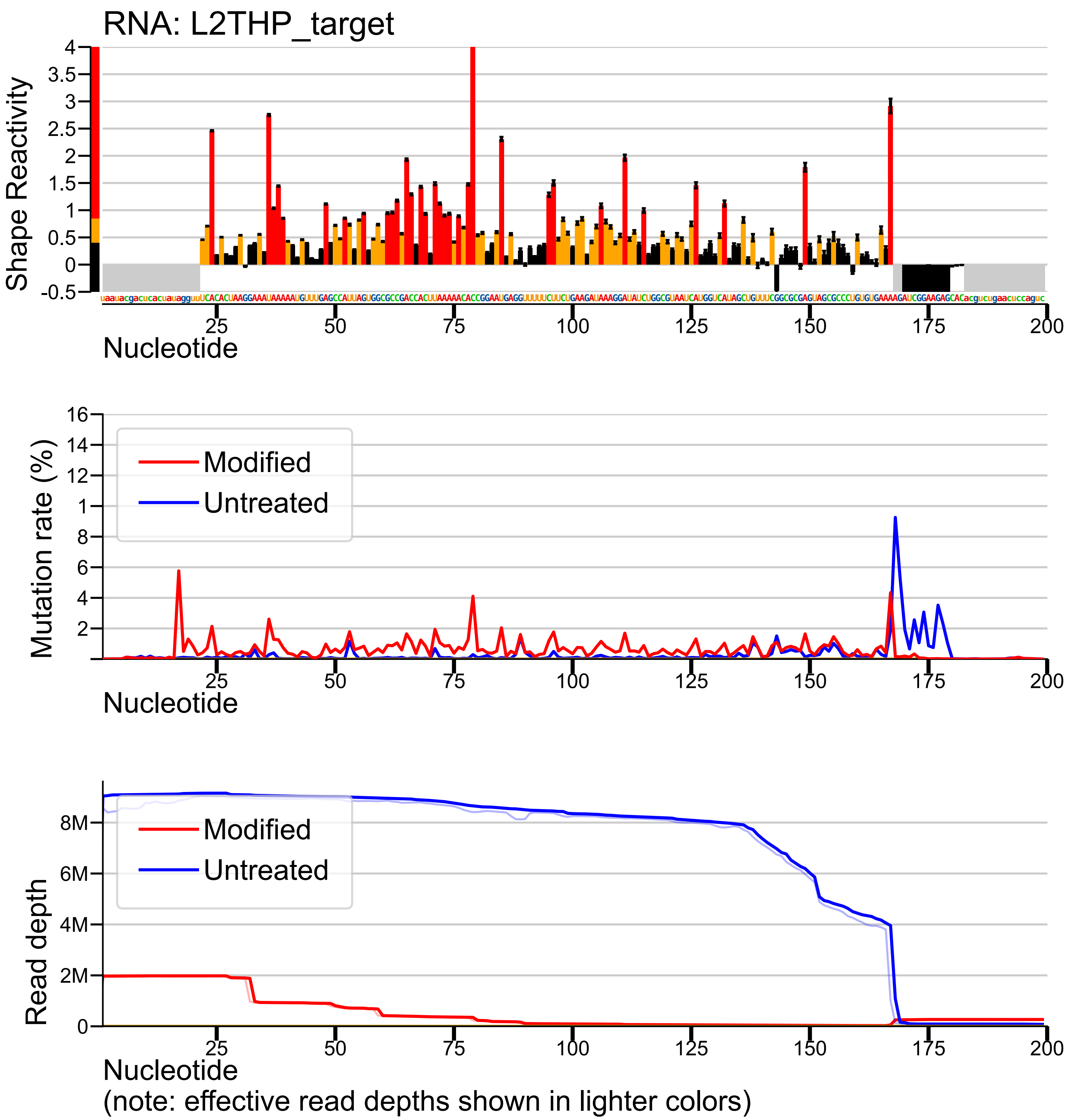
