## Supplementary material for "Ribosome-induced mRNA pseudoknot interactions visualized by DMS MaP-Seq": Target profiles 21

Note: possible data quality issue - see log file

RNA: L2THP_target

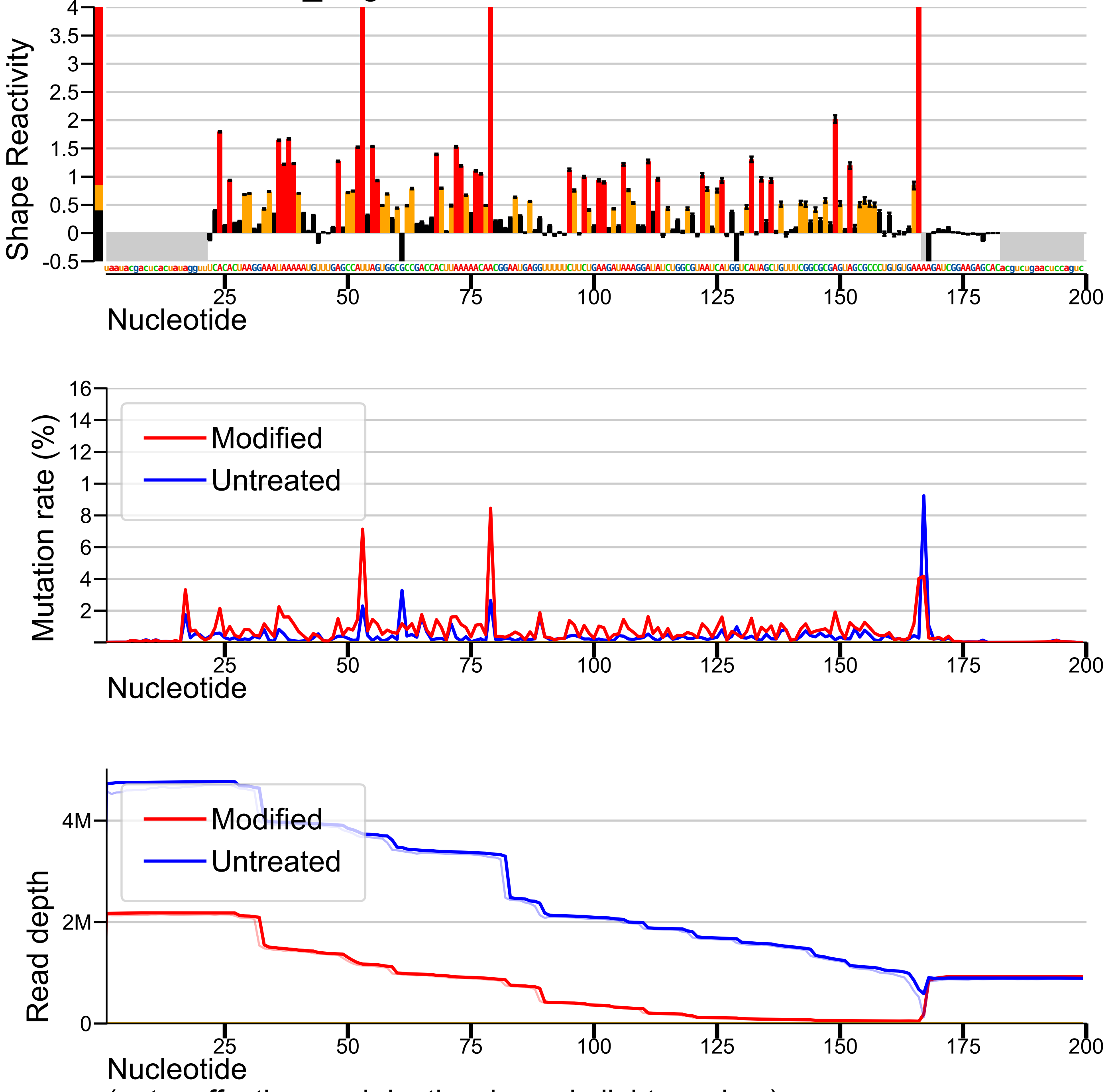


(note: effective read depths shown in lighter colors)
